# Interrogating the Regulatory Function of HAQERs during Human Cortical Development

**DOI:** 10.1101/2025.11.30.691411

**Authors:** Yashodara Abeykoon, Natalie Dzikowski, Enakshi Sinniah, Yanting Luo, Seth Weaver, Bryan J. Pavlovic, Jenelle L. Wallace, Landen Gozashti, Riley J. Mangan, Peter H. Sudmant, Craig B. Lowe, Alex A. Pollen

**Author notes:** Correspondence to Craig B. Lowe,; Alex A. Pollen. These authors contributed equally (Natalie Dzikowski, Enakshi Sinniah, Yanting Luo).

## Abstract

**Background:** Sequence divergence within gene regulatory elements has been proposed to play an important role in the evolution of human-specific traits, including cortical expansion. However, the mutational processes that efficiently modify gene regulatory elements and the target genes upon which they act are poorly understood. We investigated the regulatory function and origins of the fastest evolved regions in the human genome, termed Human Ancestor Quickly Evolved Regions (HAQERs), in their native genomic context during human cerebral cortex development.

**Results:** We identified 50 HAQERs with accessible chromatin in developing human cortex, largely arising from previously unconstrained ancestral sequences. To test the necessity of these HAQERs for gene regulation, we established an all-in-one CRISPRi lentiviral vector and linked 26 HAQERs to nearby target genes across cell types and Wnt pathway activation contexts. Rapid gains of CpGs distinguished HAQERs active during cortical development and displaying human-specific epigenomic marks. As a high density of CpG sites can drive formation of permissive chromatin, we identified 107 HAQERs with at least 17 human-specific CpG gains per kb, termed HAQER CpG Beacons. These HAQERs emerged via contributions from GC-biased gene conversion (gBGC) with evidence for selection preferentially fixing CpG sites. Notably, the *CHL1* and *DPP10* loci, both implicated in human neurological disorders, each harbored two gene-linked HAQER CpG Beacons.

**Conclusions:** Our findings reveal HAQER target genes and support a model where gBGC and natural selection jointly drive regulatory-altering CpG variants to fixation, forging regulatory innovations in the human cortex.

## Introduction

The substantial similarity in protein-coding sequences between humans and chimpanzees, has led to the hypothesis that human-specific traits are primarily driven by gene regulatory innovations [1]. Studies across diverse organisms support the role of *cis*-regulatory variation in morphological evolution [2]. Comparative genomics has revealed hotspots enriched for genetic differences that could contribute to human-specific gene regulation, including deletions (hDels) [3,4], insertions (hIns) [5], accelerated changes modifying conserved sequences (human accelerated regions, HARs) [6–8]) and accelerated changes independent of sequence conservation (human ancestor quickly evolved regions, HAQERs) [8,9].

Among hotspots of human-specific divergence, HAQERs represent the regions with the greatest number of base pair changes in the human lineage. Defined independently of sequence conservation, HAQERs encompass not only modifications of conserved elements, but also new functional elements arising from previously neutral sequences. We previously identified 1,581 HAQERs, enriched for signatures of adaptive selection [8]. The overlap of HAQERs with bivalent chromatin states and signatures of human-specific regulatory activity in the developing brain supports their role in the evolution of tissue-specific gene regulation [8,10,11]. In addition, variants in HAQERs are associated with prenatal gene expression and intracranial volume among humans and have been implicated in human language and neuropsychiatric disorders, supporting a functional role for these elements [12,13]. Reporter assays using a mouse model of neurogenesis showed that human-specific sequence changes alter the regulatory activity of several HAQERs [8], making HAQERs compelling candidate gene regulatory elements contributing to human-specific features of neurodevelopment. However, few studies have explored the target genes of HAQERs during human brain development and the sequence origins of activity differences.

In this study, we systematically examined the necessity of HAQERs in their native genomic context for regulating target gene expression during human cerebral cortex development. Since regulatory element function is highly dependent on cell types and states, we chose primary cortical progenitors that preserve the endogenous chromatin and transcriptional landscape during cortical development shaped over evolutionary timescales [14–16]. We established a primary cell culture system of human cortical radial glia, newborn neurons, and astrocytes during stages of neurogenesis predicted to influence human-specific cortical expansion [17]. By combining clustered regularly interspaced short palindromic repeat interference (CRISPRi) with single cell RNA sequencing (scRNA-seq), we identified target genes for 26 HAQERs during cortical development. Motivated by observations of Wnt-dependent changes in HAQER accessibility in cortical progenitors [18], we linked HAQERs to additional target genes in the context of Wnt stimulation. Finally, building on recent studies of species differences in enhancer activity [19], we discovered that extreme gains of CpG sites in HAQERs represents a mutational mechanism associated with human-specific gains in regulatory activity, highlighting 107 HAQER CpG Beacons. Notably, these HAQER CpG Beacons included gene-linked HAQERs regulating clinically relevant *CHL1* and *DPP10* loci that contain human-specific cortical H3K4me3 histone signatures, providing insights into the evolutionary origins and human-specific roles of these elements.

## Results

### HAQERs accessible during human cortical development emerge from previously unconstrained sequence

We sought to identify HAQERs with regulatory potential during human neurogenesis and then to measure their impact on gene expression using CRISPRi (Fig. 1a, Table S1). To identify HAQERs active during both human and non-human primate cortical development, we performed comparative assay for transposase-accessible chromatin with (ATAC) sequencing on cerebral cortex cells cultured from human and rhesus macaque (Table S1). We selected human and rhesus primary cortical progenitor cells to study HAQER function directly in matched, developmentally relevant cells with high fidelity to endogenous chromatin states and regulatory programs. Comparison with rhesus macaques, which lack the derived human-specific substitutions that define HAQERs and diverged from the human lineage roughly 25 million years ago, also informs the extent of shared regulatory activity at orthologous loci among primates. To survey regulatory potential at comparable stages of neurogenesis [17], we selected stage-matched human and rhesus mid-gestation samples according to molecular and anatomical landmarks (Fig. 1b, Fig. S1a,b) [20,21].

**Figure 1.**
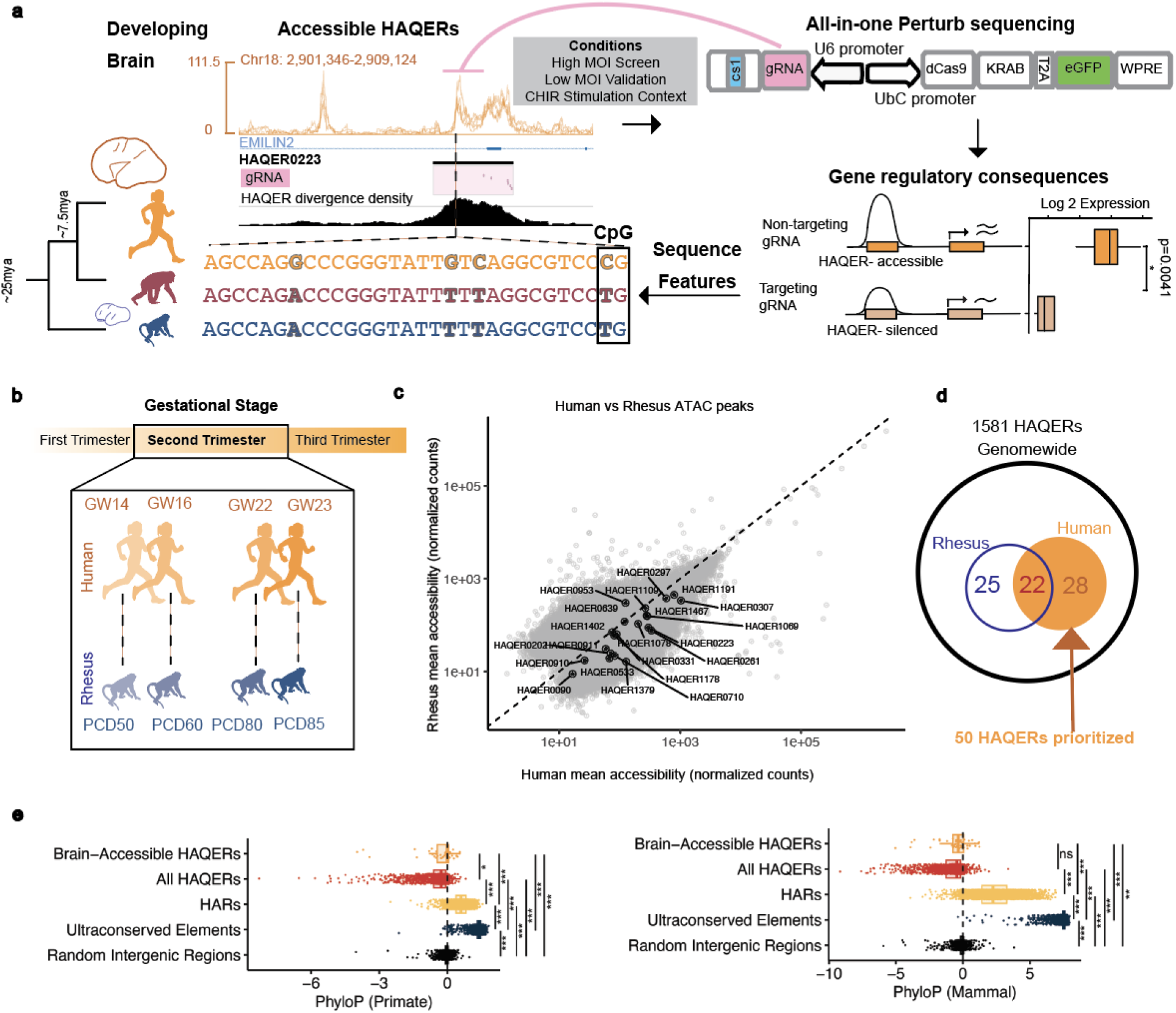
ATAC seq identifies HAQERs with accessible chromatin during human cerebral cortex development. a. Schematic of the overall study design. Accessible HAQERs in the developing cortex were identified from ATAC-seq. Shown as an example is HAQER0223 at the *EMILIN2* locus with multiple gRNAs targeting it. Comparative sequence alignment shows sequence divergence between human, chimpanzee, and rhesus over evolutionary time (mya-million years ago). Perturbation of accessible HAQERs in primary cortical progenitors enables differential expression analysis of nearby genes to interrogate HAQER regulatory function. b. Age matched human and rhesus primary cerebral cortex samples representing the second trimester of gestational stage are dissociated and plated as 2D cultures (human, n=4, rhesus, n=4). c. Scatter plot of normalized ATAC seq mean counts for X= human and Y= rhesus. d. Venn diagram showing intersection of 1581 total HAQERs with peaks detected to be significantly accessible in human vs rhesus in the second trimester developing brain. A subset of 50 HAQERs overlapping accessible chromatin in human samples prioritized to carry forward for perturb sequencing. e. Mean phyloP conservation scores from 447-way mammalian (right) and 239-way primate (left) alignments across genomic regions. Ultraconserved elements (UCEs) serve as a positive control for extreme conservation, while random intergenic regions provide a neutral baseline. While HARs show significant cross-species conservation, HAQERs exhibit slightly negative phyloP scores, consistent with elevated substitution rates across lineages[9]. (Bonferroni-corrected pairwise *t*-tests; * p<0.05; ** p<0.01; *** p<0.001; ns=not significant).Figure 2

We generated 2D cultures by first dissociating human and rhesus primary cortex tissue and then expanding the cells in media supplemented with FGF2 and EGF to enrich neural and glial progenitors [22]. Immunostaining and marker gene analysis confirmed that the majority of the cells in culture in both species were enriched for radial glia and astroglia cell types marked by GFAP (Fig. S1b,c). We performed two technical replicates of ATAC-seq for all samples and called significant peaks in each species separately. As expected, technical replicates were highly correlated and genomic loci with significantly accessible chromatin were enriched for regions nearest to transcription start sites (TSSs) in both species (Fig. S1d,e). Intersecting accessible loci in cultured cortical cells with enhancers that show reporter activity in the developing mouse (validated VISTA Enhancers) [23,24] revealed significant enrichment of brain and brain-associated enhancers compared to other tissue groups (p-value = 4.6x10^-13; Odds Ratio = 2.02) (Fig. S1f,g). Accessibility was largely correlated between orthologous peaks in humans and rhesus (Spearman’s rho = 0.790, p < 2.2e-16) (Fig.1c, Fig. S1d).

To identify HAQERs accessible in developing cortex, we intersected 1,581 HAQERs with ATAC peaks detected to be significantly accessible in human and rhesus samples. We identified 50 HAQERs that overlapped with accessible peaks in at least one human developmental stage (Fig. 1d), with 22 of these HAQERs also being accessible in rhesus, and an additional 25 HAQERs only reaching significance in rhesus. HAQERs showed correlated chromatin accessibility in human and rhesus samples (Spearman’s rho = 0.776, p < 2.2e-16), indicating that these human-specific mutations likely arose in loci with ancestral chromatin accessibility (Fig. 1c). Differential accessibility analysis of orthologous peaks confirmed that HAQERs classified as accessible in only one species had higher accessibility in that species, whereas HAQERs accessible in both species, including HAQER0297 in the CHL1 promoter region, generally showed higher accessibility in human than in rhesus samples Fig. S1h). Despite evidence for ancestral accessibility in many cases, brain accessible HAQERs, similar to all HAQERs, showed limited sequence constraint compared to HARs and ultraconserved elements, across mammals and primates, indicating that brain accessible HAQERs did not experience sustained negative selection over macroevolutionary timescales (Fig. 1e). Of the 50 HAQERs accessible in human samples, 30 were within 500 base pairs (bp) of the TSS of the nearest gene, and termed proximal elements, while the remaining 20 were further than 500 bp from a nearest TSS and were termed distal elements (Table S2). We hypothesized that epigenetically silencing these 50 HAQERs displaying accessible chromatin in our samples would result in altered expression of the genes they regulate during human cerebral cortex development.

### HAQERs regulate gene expression in human cortical development

While previous high-throughput enhancer assays identified HAQERs with sufficient regulatory activity to drive reporter gene expression in mouse embryonic brain [8], the target genes of HAQERs in their native genomic context during human brain development remain unclear. Addressing this gap, we epigenetically repressed 50 HAQERs displaying chromatin accessibility in cortical progenitors using CRISPRi and measured transcriptional consequences using scRNAseq (Perturb-seq). To account for biological and technical variation, we performed experiments across four human samples and used a multiplexed pool of single guide RNAs (sgRNAs) to target all HAQER candidates in shared cell culture conditions (Fig. 2a, Table S3).

**Figure 2.**
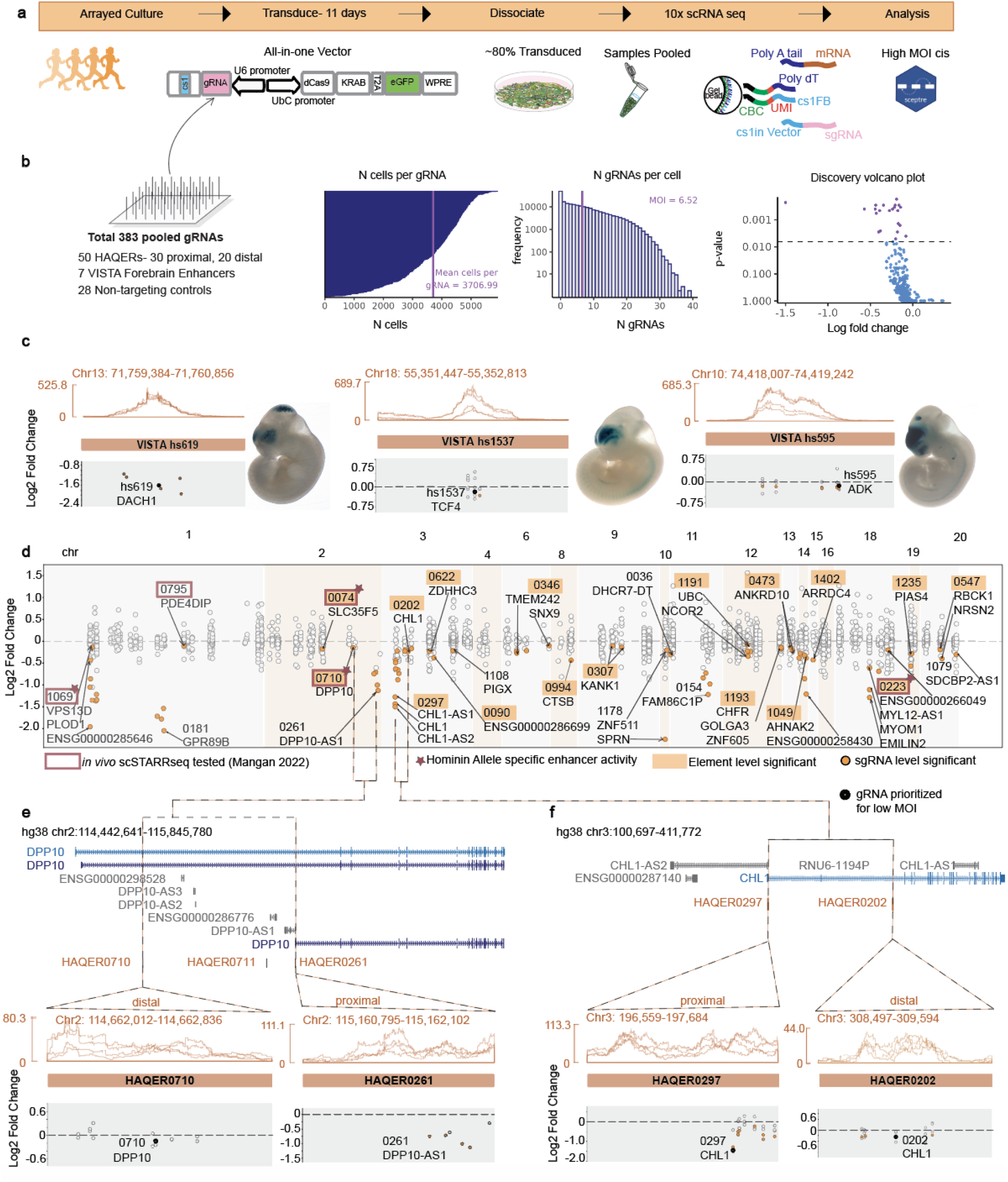
High MOI Perturb-seq links HAQERs to their *cis*-regulated genes in developing human cortex. a. Schematic of experimental design for transducing 4 human samples with the all-in-one direct capture Perturb-seq lentiviral vector. b. Left to right: Schematic of pooled sgRNA oligonucleotides cloned to the vector. Output statistics from SCEPTRE high MOI cis analysis showing cells/gRNA, gRNAs/cell (MOI), significant regulatory element-gene pairs identified at the SCEPTRE element level analysis. c. Chromatin accessibility of VISTA forebrain enhancer elements included in high MOI screen to increase confidence (top). relative location of sgRNA and the log_2_FC change of its paired cis gene plotted along the VISTA element genomic locus (bottom). d. HAQER paired genes with SCEPTRE high MOI analysis plotted along their relative genomic locus. log_2_FC of paired gene expression in cells with target sgRNA compared to cells without that specific gRNA is plotted in Y axis. Each X axis location of a dot is the relative position of the sgRNA coordinates in the hg38 genome. Each HAQER was targeted with multiple sgRNAs. The HAQER-gene pair dot with the highest significant log_2_FC is labeled in the plot with a black arrow. Whenever multiple genes were linked to a HAQER, each of the pairings with the highest fold change levels were labeled. The vertical alignment of dots along the Y axis represent all the genes in a window of 0.5Mb tested by SCEPTRE for each sgRNA. Only the significant pairings are colored as orange dots while non-significant pairings are in white. HAQERs previously tested in *in vivo* scSTARRseq are boxed in red and labeled with an additional star symbol if they were reported to elicit human-specific enhancer activity[8]. e. Magnified view into HAQERs regulating *DPP10* and *CHL1* genes. From top to bottom: UCSC hg38 genome browser view of *DPP10* and *CHL1* genes. Each gene contains a distal and a proximal HAQER that show accessible chromatin. Relative position of sgRNAs targeting each HAQER and log_2_FC of the HAQER paired cis gene.

To support analysis in primary cells and mitigate the effects of viral recombination on multiplexed pools, we created an all-in-one lentiviral vector from a pre-existing CRISPRi vector [25] by inserting a capture sequence (cs1) to the sgRNA scaffold, enabling repression of HAQERs in unmodified cell lines and direct sequencing of sgRNAs in each cell (Fig. S2a). To assess knockdown efficiency, we transduced primary human cells with the lentiviral all-in-one vector carrying an sgRNA targeting the Beta-2-microglobulin (B2M) surface protein encoding gene. Compared to cells which received the non-targeting control, cells which received the B2M-targeting sgRNA showed >90% knockdown of B2M protein quantified by flow cytometry (Fig. S2b, Methods).

After confirming the efficiency of our Perturb-sequencing platform in primary cortical progenitor cell culture, we cloned a pool of 383 unique sgRNAs into the all-in-one vector with 3-7 individual sgRNAs targeting each of the 50 HAQERs (Fig. 2b). This sgRNA pool also included 7 VISTA [23,26] forebrain enhancer elements (hs111, hs200, hs595, hs619, hs646, hs724, and hs1537) as controls. The VISTA forebrain enhancers were not true positive controls, since their necessity for *cis*-regulation of target genes had not been analyzed before. However, they provided candidate enhancer elements to assess the sensitivity in the Perturb-seq assay analysis since their sufficiency to drive significant LacZ expression in developing mouse brains is well established. We included 28 previously published non-targeting sgRNA sequences as our negative controls ([27]).

We transduced the human progenitors at developmental stages gestational week (GW)14, 16, 22 and 23, with the sgRNA library targeting HAQERs, VISTA elements, and non-targeting controls cloned into the lentiviral all-in-one Perturb-seq vector. To efficiently assess noncoding element function, we first performed high multiplicity of infection (MOI) screening, in which cells are transduced with multiple sgRNAs [28]. After 11 days of transduction, we captured single cells with 10x 3’ HT RNA-seq technology. Since the cs1 was inserted directly in the sgRNA scaffold of the vector carrying each sgRNA, we were able to directly capture the gene expression profile and the sgRNA in each cell together. We observed a mean MOI of ∼7 sgRNAs per cell (Fig. 2b). For each of the 50 HAQER elements and 7 VISTA elements, we analyzed downregulation of genes located within a 0.5 Mb window. After quality control for the expression level of nearby genes and number of cells expressing the sgRNA, we considered 348 candidate gene linkages to these 57 elements using SCEPTRE, a well-calibrated statistical method for connecting perturbations to molecular phenotypes that controls for multiple hypotheses ([29,30]).

SCEPTRE considers the collective impact of each sgRNA targeting the same element to evaluate gene expression effects at the regulatory element level [29]. CRISPRi-mediated repression of 22 regulatory elements significantly downregulated the expression level of at least one candidate *cis*-target gene within 0.5 Mb at a Benjamini-Hochberg (BH) FDR of 10% (Fig. 2b, Fig. S2c, Table S5). These 22 regulatory element-gene linkages included 5 of 7 VISTA forebrain enhancer elements, hs111, hs595, hs619, hs646, and hs1537 paired with significant downregulation of *GLI3*, *ADK/KAT6B*, *DACH1*, *METAP1D*, and *TCF4*, respectively (Fig. 2c, Table S5). Notably, the effects on target genes in these control enhancers varied, with hs619 showing strong effects for all sgRNAs (median log_2_fold change (FC) = -1.471), while other sgRNA-gene pairs showed more modest effects in the range of log_2_FC -0.139 to -0.394 and a lower fraction of active sgRNAs more typical of CRISPR studies of enhancers [31,32]. Most of the control enhancers regulated a single candidate *cis*-target gene, but targeting hs595 reduced the expression of both *ADK* (3/5 sgRNAs with log_2_FC < -0.195) and *KAT6B* (5/5 sgRNAs with log_2_FC < -0.250). Among HAQERs, targeting 17 out of 50 accessible elements significantly reduced the expression of at least one gene within 0.5 Mb in the element-level analysis (Fig. 2d, orange boxes).

Since each regulatory element was targeted with 3-7 sgRNAs per element, we next focused on differential expression at the gRNA level. For this we performed SCEPTRE analysis for each unique sgRNA with a reduced cell number threshold. This analysis showed 26 HAQERs paired with significant downregulation of candidate *cis*-target genes at a 10% FDR (Fig. 2d, Table S6), nominating 9 additional candidates. Among the HAQERs significantly downregulating target genes, HAQERs 0074, 0223, 0710, 0795, 1069, had previously been tested for enhancer activity with *in vivo* scSTARR-seq [8]. This previous experimental comparison of derived and ancestral alleles revealed that HAQERs 0074, 0223, 0710, 1069 showed hominin sequence-specific increases in enhancer activity (Fig. 2d). While most HAQERs regulated at least one target gene, we noticed several examples of HAQERs linked to multiple genes, long non-coding RNAs, and/or sense-antisense transcripts of the same gene body. Of note, we observed that *CHL1* and *DPP10,* genomic loci associated with neurological disorders, each harbored multiple HAQERs (Fig. 2e,f). The *DPP10* gene body harbored HAQERs 0261, 0710, and 0711. HAQERs 0261 and 0710 showed chromatin accessibility in our samples and were targeted with multiple sgRNAs to test their regulatory function. Silencing HAQER0261 showed significant reduction in its proximal transcript *DPP10-AS1* while HAQER0710 showed reduction in its distal *DPP10* sense transcript (Fig. 2e). In the *CHL1* locus, both proximal HAQER0297 and distal HAQER0202 were linked to downregulation of *CHL1* (Fig. 2f), with the repression of the proximal element driving a stronger reduction in the target gene. Together, high-throughput CRISPRi screens nominated multiple HAQERs as functional regulators of gene expression during cortical development.

### Reproducibility and cell type-specificity of HAQER regulatory function

We next sought to validate the observed HAQER-gene linkages with a higher resolution and statistical power using low MOI Perturb-seq (Fig. 3a, Fig. S3a). Our primary cell cultures were enriched for radial glia and astroglia lineages, but oligodendrocyte (oligo), inhibitory and excitatory neuron lineages were also present at lower abundance (Fig. 3b, Fig. S2d,e, Fig. S3b). This heterogeneity enabled us to further investigate the cell type specificity of HAQER-gene linkages. For each significant HAQER, we prioritized the 1-2 sgRNAs with the strongest activity in the high MOI screen, resulting in a total of 29 sgRNAs targeting 17 HAQERs, 4 VISTA forebrain enhancers, and 4 NTCs. We cloned the 29 sgRNA library into the all-in-one lentiviral vector, transduced the same four human samples at a low MOI, enriched for GFP-expressing cells by fluorescence activated cell sorting (FACS), and performed differential *cis* gene expression analysis (Fig. 3a, Fig. S3c, Table S7). The effects of sgRNAs on candidate target genes were strongly correlated between high and low MOI experiments (R^2^=0.745) (Fig. 3c, Table S8).

**Figure 3.**
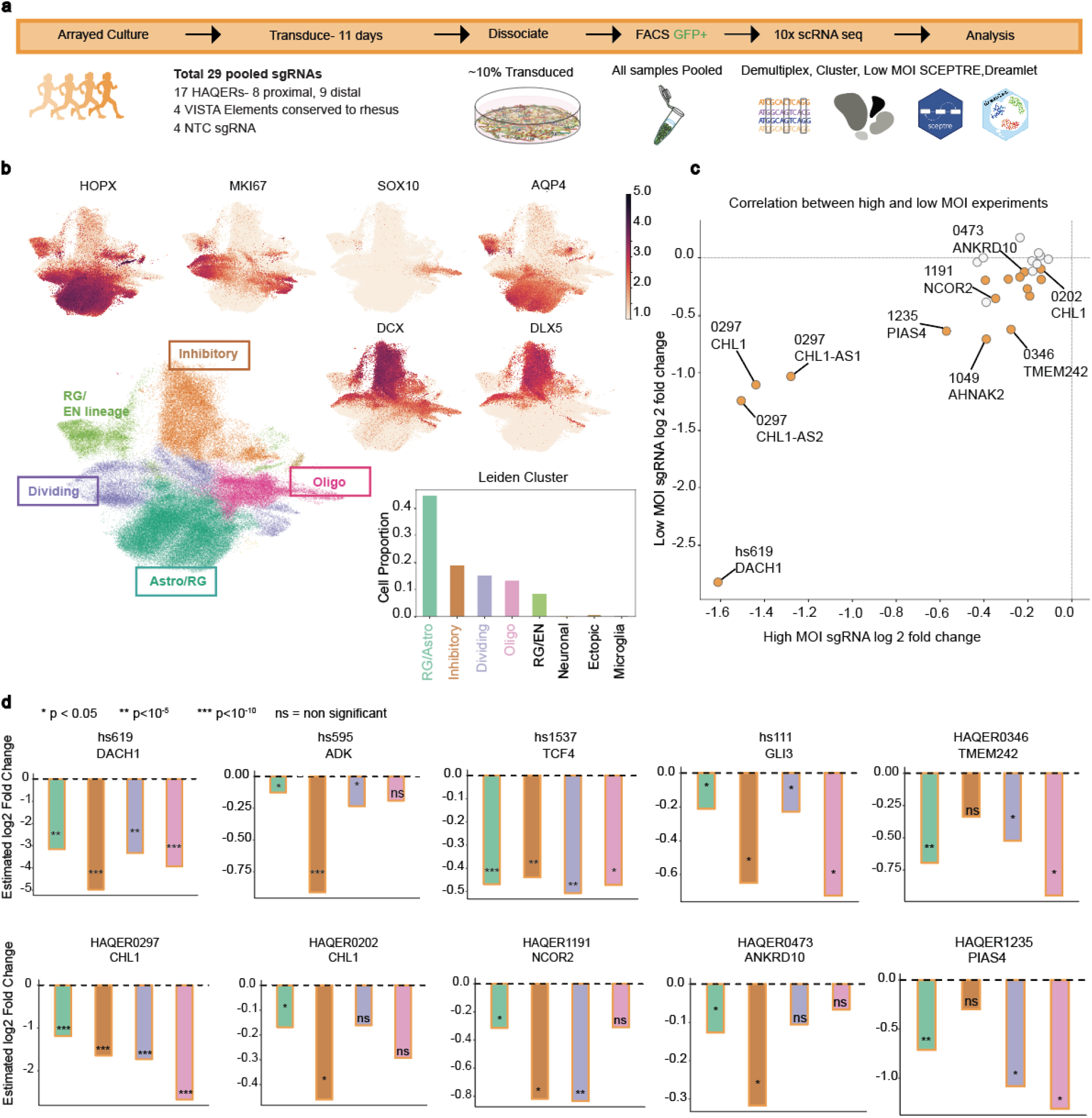
Low MOI Perturb-seq reproduces the gene regulatory function, and shows brain cell type-specificity of HAQERs. a. Schematic of experimental design for low MOI Perturb-seq for prioritized HAQERs from initial high MOI screen. b. UMAP of 112,846 single cells colored by Leiden clusters and annotated by reference mapping and normalized expression of marker genes *HOPX, MKI67, SOX10, AQP4, DCX*, and *DLX5*. The bottom right shows the proportion of cells in each cluster. c. Correlation of HAQER-gene pairs identified from the previous high MOI screen with the low MOI validation. Each dot represents a specific sgRNA-gene pair tested for linkage (Log_2_FC for targeting-sgRNA vs NTC). Colored dots represent significance. d. Log_2_FC of the HAQER paired gene in different cell clusters assigned to Astro/Radial Glia, Inhibitory, Dividing, and Oligodendrocyte lineages.

As enhancers are frequently cell type-specific [33], we next examined HAQER and VISTA element linkages to their target genes across cortical cell populations. For statistical power in the differential gene expression analysis, we focused on cell clusters RG/Astro, inhibitory, dividing, and oligo that had the highest number of cells captured (Fig. 3b). Targeting VISTA forebrain enhancers reduced the expression of paired genes in all four cell types with varying degrees of cell type specificity in effect sizes, notably for hs595 which showed stronger effects in inhibitory neurons than the other cell types (Fig. 3d). Similarly, targeting proximal HAQER0297 downregulated *CHL1* in all four cell types. In contrast, targeting distal HAQERs 0202, 1191, 0346, 0473, 1235, and 1049 drove cell type-specific downregulation of respective paired genes *CHL1*, *NCOR2*, *TMEM242*, *ANKRD10*, *PIAS4*, and *AHNAK2* (Fig. 3d). In each case, expression of candidate *cis*-target genes was reduced in the radial/astro glia cluster, but in several cases we observed strong downregulation in other cell types, including in inhibitory neurons for the HAQER0202-*CHL1* linkage (Fig. 3d).

In parallel, we generated data from age-matched rhesus macaque samples cultured and processed under identical experimental conditions (Fig. S3a,d). The sgRNA protospacer and the protospacer adjacent motif (PAM) sequences targeting HAQERs showed limited conservation between human (hg38) and rhesus (rheMac10) genome assemblies due to the rapid sequence changes in the human lineage. This divergence enabled rhesus data to serve as an additional layer of negative control to further rule out off-target effects of sgRNAs in our library. For example, targeting HAQER0622 elicited strong transcriptional responses with a distinct cluster formed exclusively by a subset of cells expressing the sgRNA (Fig. S3e). However, in rhesus, the sgRNA had 3 mismatches for the homologous site including a mismatch 3bp away from the PAM site, but still resulted in a homologous ectopic cluster (Fig. S3e), arguing that other effects of this sgRNA, independent of the human-specific target site, drove the downstream transcriptional changes. While sequence divergence in HAQERs limited direct comparison with rhesus orthologs, sgRNAs targeting HAQERs 1191 and 0473 showed only 1bp divergence between human and rhesus in the target specific crRNA (Fig. S3f) and elicited stronger gene repression in human than rhesus (Fig. S3f,h). Conversely, targeting VISTA elements in rhesus led to downregulation of the same paired genes we observed in human samples (Fig. S3g,h). Together, the concordance between low and high MOI CRISPRi experiments and cell type level analysis support the necessity of the HAQERs for regulating target genes during cortical development.

### Wnt stimulus-dependent HAQER-gene linkages

Wnt signalling influences cortical development and patterning as a major morphogen and mitogen, with elevated activity driving increased cell proliferation [34] and altered signaling linked to neurodevelopmental disorders [35]. Stimulating Wnt signaling by adding Wnt3a or inhibiting GSK-3, a downstream kinase that suppresses Wnt pathway activation, alters the accessibility of thousands of regulatory elements and genes in cultured primary cortical progenitors [18], implying that stimulation may alter regulatory element gene linkages. The overlap between 1581 HAQERs and accessible chromatin in cortical progenitors increased from 23 at baseline to 37 following Wnt stimulation in a recent study [18]. Of the HAQERs linked to genes at the element level, 9 out of 17 showed accessible chromatin following Wnt pathway stimulation, including 3 HAQERs 0202, 0710, and 1191, that were exclusively detected as accessible under Wnt stimulation condition [18]. This overlap motivated us to inspect the regulatory potential of HAQERs under Wnt pathway stimulation.

We transduced human and rhesus cells with the same 29 sgRNAs as in the validation assay including HAQER0202, 0710, and 1191 targets which show Wnt dependent increased chromatin accessibility [18]. For this experiment, we additionally stimulated cells on post-transduction day 9 with 2.5 µM CHIR (CHIR99021), a selective GSK-3 inhibitor, for 48 hours (Fig. 4a). We then captured single cells 11 days post transduction using 10x scRNA-seq. Visualization of genes previously noted to be differentially expressed in developing brain cells during CHIR stimulation, such as increased *AXIN2* and decreased *TCF7L2*, indicated a response to Wnt signaling [18] (Fig. 4a-d).

**Figure 4.**
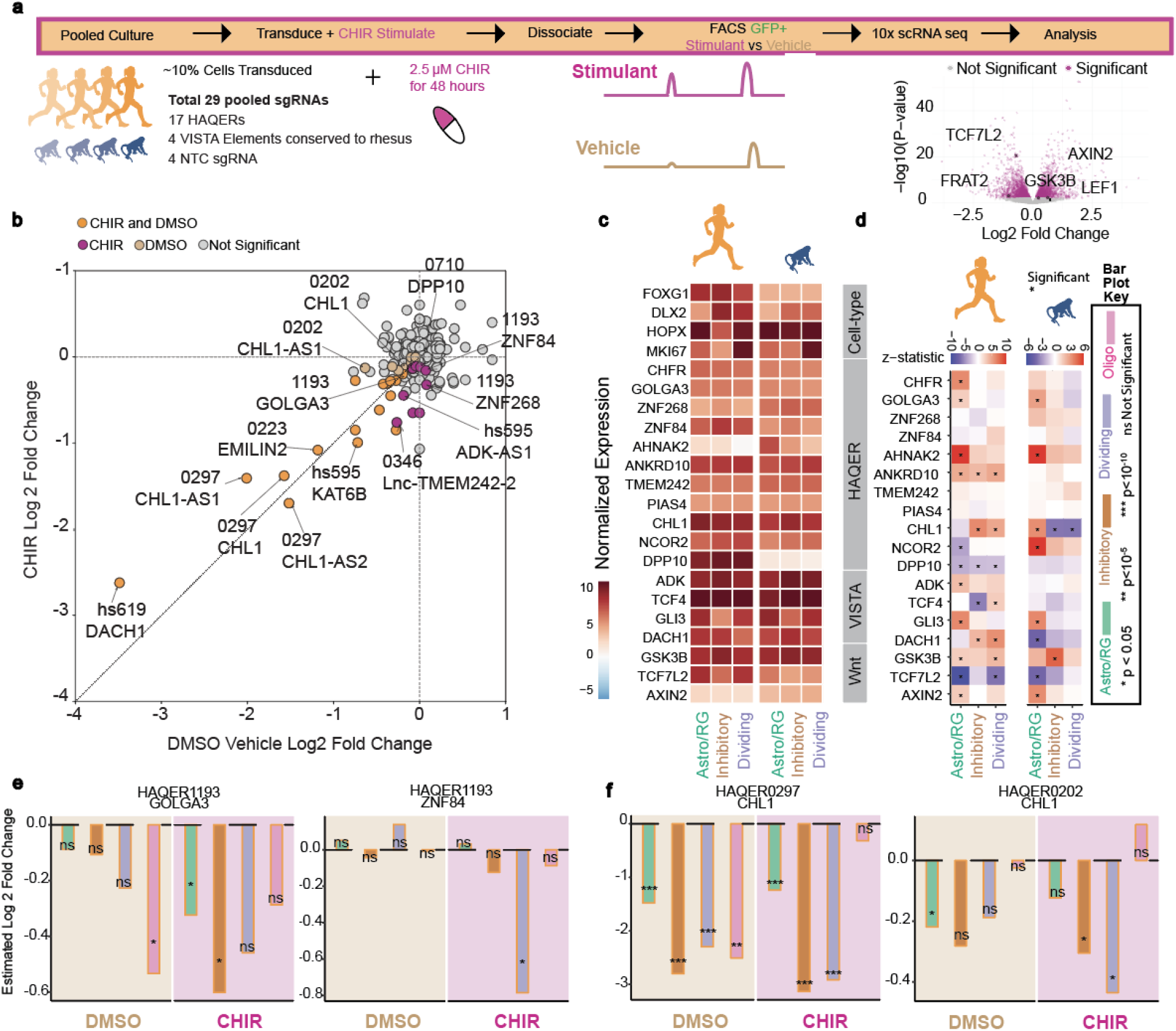
CHIR stimulation context influences HAQER gene regulatory effects in developing cortex. a. Schematic of the study design. Cells are transduced at an MOI of 1 sgRNA per cell and CHIR stimulated prior to single cell capture. Schematic of differential chromatin accessibility dependent on the context. Volcano plot showing differential gene expression of cells transduced with NTC exposed to CHIR stimulation compared to DMSO vehicle. b. Scatter plot of HAQER-gene linkages in DMSO Baseline (X axis) vs under CHIR stimulation (Y axis) for human samples. Each dot represents a HAQER-gene linkage (log2FC for targeting-sgRNA vs NTC in each condition). Colors of the dots represent significance of the linkage in baseline, CHIR, both, or none. c. Heatmap of normalized expression of genes linked with Wnt pathway, VISTA, HAQER and cell-type markers, in cells containing NTC sgRNA in the DMSO baseline condition. Human and rhesus panels displayed side by side. d. CHIR responsive expression level changes of Wnt, VISTA, and HAQER-linked genes in human and rhesus samples. Z-statistic notes the direction of the gene expression change compared to DMSO. e. Cell type-specific gene expression level changes of HAQER-linked genes in the context of DMSO or CHIR.

When analyzing all cell types combined, the effects of targeting HAQERs on paired genes were highly correlated between DMSO Baseline (Table S9) and CHIR stimulation (Table S10) conditions (R^2^= 0.723) (Fig. 4b). Targeting VISTA positive control elements showed highly correlated paired genes and effect sizes in both combined and cell-type specific analyses between CHIR and DMSO (Fig. 4b, Fig.S4a). However, we observed differences in paired gene expression as well as cell type specificity for a subset of HAQER perturbations under CHIR stimulation compared to DMSO baseline. For example, HAQER1193 located in an out of frame exon of the *CHFR* gene paired with *GOLGA3* downregulation under both baseline and CHIR stimulation contexts. In contrast, HAQER1193 paired with additional nearby zinc finger genes in a context dependent manner (Fig. 4e, Fig. S4a,b). HAQERs 0297 and 0202 in the *CHL1* locus responded to CHIR stimulation in cell type specific ways. Proximal HAQER0297 paired with downregulation of *CHL1* under both CHIR stimulation and baseline. Compared to 0297, distal HAQER0202 showed more cell type specificity under different contexts. Targeting HAQER0202 under baseline conditions reduced *CHL1* primarily in the RG/Astro cell cluster, while targeting under CHIR stimulation reduced *CHL1* in inhibitory and dividing cell clusters (Fig. 4f). HAQER0710 paired with *DPP10* under CHIR stimulation when all cell types were combined, but not in cluster level analysis where we had lower statistical power (Fig. 4b). Visualization of Genehancer tracks [26,36] in the UCSC genome browser [37] for *CHFR*, *CHL1*, and *DPP10* genetic loci showed direct and/or indirect looping between the above HAQERs and their linked genes, including under stimulation conditions, supporting observed HAQER-gene linkages (Fig. S4b,c,d).

Additionally, we investigated the HAQER, VISTA enhancer, and Wnt pathway-linked gene expression levels at baseline (Fig. 4c) and their changes under CHIR (Fig. 4d), in cells containing NTC sgRNAs in human vs. rhesus. These genes in the DMSO baseline had broadly comparable expression levels between human and rhesus (Fig. 4c). In concordance with prior studies, we observed that the *DPP10* gene linked with HAQERs 0710 and 0261 in our study was expressed at a higher level in the human samples compared to rhesus [38] (Fig. 4c). A subset of HAQER-linked genes showed CHIR responsive significant increase or decrease in expression levels in human and rhesus. For example, between the two species, the direction of CHIR response was similar for HAQER-linked genes *CHFR, GOLGA3,* and *AHNAK2*. Conversely, *NCOR2* and *CHL1* genes showed opposite responses between human and rhesus under CHIR stimulation, suggesting a species-specific transcriptional response to CHIR in these 2 HAQER-linked genes (Fig. 4d). Collectively, these findings show a largely conserved landscape of HAQER-gene linkages following Wnt stimulation, while revealing additional HAQER gene linkages and changes in cell type specificity.

### HAQER sequence variation overlaps with human-specific H3K4me3 histone and CpG dinucleotide signatures

We next examined the human-specific regulatory properties and underlying sequence changes of HAQERs accessible in cortical progenitors and those linked to genes in our high-MOI Perturb-seq assay (Fig. 5a). We overlapped 50 accessible HAQERs, and the gene-linked HAQERs, with previously published datasets of human-specific epigenetic signatures. HAQERs accessible during human cortical development, including gene-linked HAQERs, showed significant overlap with human-specific H3K4me3 peaks in the prefrontal cortex compared to chimpanzee and rhesus [38] (17/50 accessible in developing cortex, 5/17 element level gene-linked, 8/26 sgRNA level gene-linked; n=10,000 permutations, p < 1×10⁻⁴ for all, Table S11, Fig. 5b), suggesting HAQER sequence divergence contributes to human-specific epigenetic signatures.

**Figure 5:**
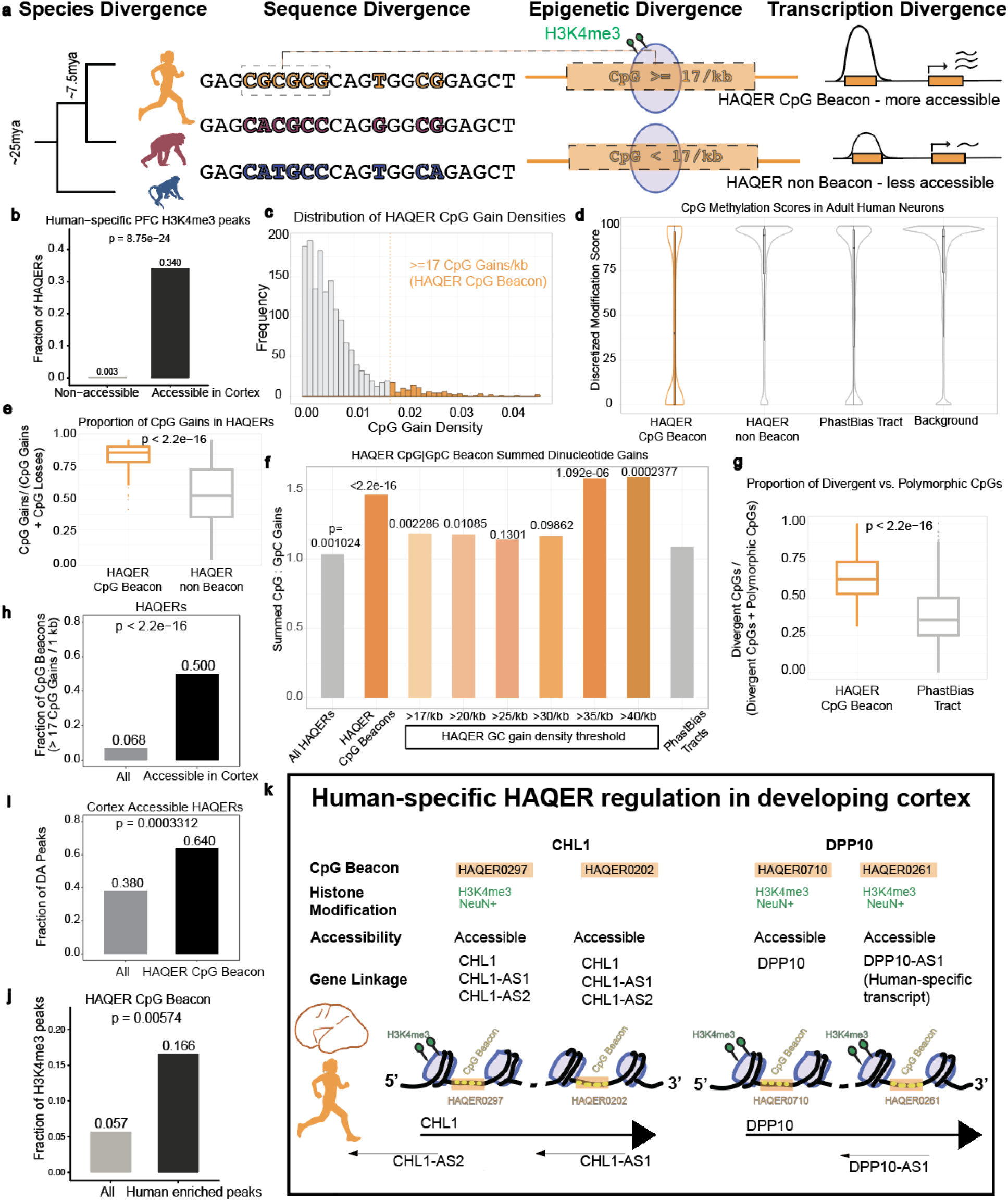
Human-specific epigenetic signatures overlapping HAQER CpG Beacons. a. Cartoon of HAQER sequence divergence overlapping with human-specific epigenetic and transcription divergence, emphasized by accessible and gene-linked HAQERs from our study. Species divergence shows the evolutionary tree of human, chimpanzee, and rhesus. HAQER sequence divergence with SNPs between human and non-human primates colored for each species (human-orange, chimpanzee-magenta, rhesus-blue). Epigenetic divergence showing dense CpG clusters leading to deposition of H3K4me3 marks in green on histones in purple. Proposed model of genetic and epigenetic modifications of HAQERs leading to functional differences in their accessibility and expression changes in genes targeted by these HAQERs. b. Bar chart analyzing previously published H3K4me3 ChIP-seq signal in postnatal human vs NHP PFC [38] (chimpanzee and rhesus). Accessible HAQERs in developing human cortex from our study are significantly enriched for human-specific H3K4me3 signatures compared to non-accessible HAQERs. c. Frequency histogram of HAQER CpG gain density. Orange dotted line marking HAQER CpG beacons thresholded at >= 17/kb CpG density. d. Violin plot of DNA methylation scores at HAQER CpG Beacons (orange) versus non-Beacon HAQERs, phastBias tracts, and background regions (grey) in adult human brain neurons, pooled across two individual donors (ages 60 and 84 years). HAQER CpG Beacons show significantly reduced DNA methylation compared to all other genomic regions (Mann-Whitney U test, p < 2.2 × 10⁻¹⁶ for all pairwise comparisons; pairwise test statistics in Table S15). e. Boxplot showing proportion of CpG gains compared to losses in HAQER CpG Beacons (>= 17/kb) vs non-Beacons. f. Bar chart for the ratio of summed CpG over GpC gains (y axis) in different subsets of HAQERs crossing varying thresholds for CpG gains or GpC gains. A HAQER is included in the subset if it crosses the threshold due to either CpG gains or GpC gains, with the two types of gains considered separately. Significant p value calculated vs phastBias tracts (regions of biased GC fixation genomewide) not overlapping a HAQER CpG Beacon. From left to right bars reflect GC gain for sets of: all HAQERs (N=8851/8600), HAQER CpG Beacons (2206/1511), HAQERs with CpG or GpC gain densities of >17/kb (2456/2079), >20/kb (1910/1629), >25/kb (1241/1092), >30/kb (777/669), >35/kb (410/260), >40/kb (219/138), phastBias tracts (48898/45150) g. CpG site Divergence vs Polymorphism for HAQER CpG Beacons vs phasBias tracts. phastBias tracts overlapping a HAQER CpG Beacon or with zero counts were excluded. h. Bar chart showing increased accessibility of the HAQER CpG Beacons compared to the set of all HAQERs in the developing human cortex. i. Bar chart of HAQER CpG Beacons showing increased differential accessibility between human and rhesus when compared to all HAQERs. j. Bar chart showing HAQER CpG Beacons are significantly overrepresented among human-enriched H3K4me3 peaks in the developing cortex, compared to all HAQERs, assayed by H3K4me3 CUT&Tag in human and rhesus k. Cartoon of levels of regulation involved with HAQER divergence in at *CHL1* and *DPP10* genomic loci. HAQER sequence divergence contains an enrichment for the gain of CpGs in human lineage. HAQERs 0297, 0202, 0710, and 0261 are CpG Beacons (>=17 CpGs/ 1kb) and HAQERs 0297, 0710, and 0261 show human-specific enrichment of H3K4me3 histone modifications in NeuN+ cells in PFC. HAQERs at the *CHL1* locus regulate both sense and anti-sense transcripts. HAQERs at the *DPP10* locus regulate sense or anti-sense transcripts in a mutually exclusive manner in the developing human cortex.

One proposed mechanism for the gain of H3K4me3 involves the formation of hypomethylated clusters of CpGs [19,39]. These sites bind CFP1, which in turn recruits chromatin modifying enzymes, including SETD1 that methylates H3K4 [40,41]. Motivated by this process as a source of regulatory innovation in the human lineage, Bell and colleagues identified 21 loci, termed extreme CpG beacon clusters, with at least 20 human-specific CpG sites within 1kb [42] and showed their significant overlap with the set of human-specific H3K4me3 signatures [38,39], which are the same human-specific signatures that largely overlapped with the accessible and gene-linked HAQERs [38,39]. Strikingly, 13 of these 21 extreme CpG beacon clusters overlapped with HAQERs, including 3/17 gene-linked HAQERs (0223, 0297, 0710), and HAR1 (also identified as HAQER0035) [6] (Fig. S5a, Table S12).

To better understand the extent to which high-density CpG gains contributed to HAQER formation, we revisited the calculation of CpG Beacon clusters with new data and refined methods. Advances over the last decade allowed us to use an alignment including the bonobo genome, improved assemblies for all great ape species (including humans), and an explicit inference of the human-chimpanzee ancestral state [8]. In a follow-up publication, authors from the original CpG Beacon paper published a refined threshold of 17 or greater human-specific CpG gains per kb [39], which we now used to analyze each of the 1,581 total HAQERs. At this refined threshold, 107 HAQERs were enriched for CpG beacon clusters (HAQER CpG Beacons) (Fig.5c, Table S13). To increase confidence in this threshold, we additionally assigned a significance to each HAQER based on a deviation from the genome-wide rate of CpG formation (Methods), with 102 of the 107 also showing genome-wide significance with this secondary method (p < 0.01) (Fig. S5a).

We hypothesized that these HAQER CpG Beacons would show signatures of hypomethylation in a neuronal cell type, enabling them to bind CFP1 and receive human-specific epigenetic modifications. To test this hypothesis, we conducted methylation-sensitive long-read sequencing of neurons from two individuals. HAQER CpG Beacons showed significant DNA hypomethylation compared to other HAQERs, phastBias tracts, and genomic background sequences, (Fig. 5d). Additionally, we examined non-neuronal cell types, oligodendrocytes and sperm and these cell types also showed HAQER CpG Beacons to be significantly hypomethylated (Fig. S5b). Finally, analysis of publicly available ChIP-seq data for CFP1 revealed a significant overlap between CFP1-bound DNA and HAQERs CpG beacons compared to other HAQERs and phastBias tracts (Fig. S5c). These findings nominate the gain of CpG sites, DNA hypomethylation, and chromatin modifications as a molecular cascade that links the sequence divergence of HAQERs to the emergence of human-specific regulatory function in the developing brain.

### GC-biased gene conversion plays a role in forging regulatory elements in the human genome

While the majority of HAQERs showed background rates of CpG gain, the 107 HAQER CpG Beacons represented a unique subset of HAQERs, marked by a tendency for CpG gains (Fig. 5e). If this gain was driven solely by rapid sequence change or turnover, we might expect HAQER CpG Beacons to show a similarly elevated tendency for CpG losses. However, the set of HAQER CpG Beacons showed a strong bias for the gain of CpGs, without a matched tendency to lose these dinucleotides (Fig. 5e,f), consistent with CpG gain in these regions being a directed process.

GC-biased gene conversion (gBGC) [43], a recombination-driven evolutionary process that preferentially fixes mismatches to G/C over A/T alleles, could provide a molecular mechanism underlying the directed CpG gain in HAQER CpG Beacons. Consistent with this model, 9 out of the top 10 regions of clustered GC-biased substitutions in the human genome (Table S14) [44] overlapped with HAQER CpG Beacons. The contribution of gBGC was further reinforced by the significant enrichment (p < 10^-79^) of HAQER CpG Beacons in phastBias tracts, which mark regions of biased GC fixation genomewide [45] (Fig. S5d). The chromosomal location of HAQER CpG Beacons is also consistent with gBGC due to their enrichment in telomeres and subtelomeres (Fig. S5d) [44,45]. Taken together, these findings support gBGC as an evolutionary mechanism underlying the rapid gain of CpGs in these 107 HAQERs.

GC-biased gene conversion can occur both when there are weak-strong mismatches between alleles at a given locus (interallelic) and when there are weak-strong mismatches between paralogous loci (interlocus) [43]. While both types of gBGC can result in a bias towards weak-to-strong substitutions at a site, there are also differences. Interallelic gBGC can mimic selection where a variant already present in the population is pushed to higher frequency. Interlocus gBGC can elevate the mutation rate at a site, by introducing G/C variants already present at a paralog elsewhere in the genome. Interallelic gBGC is likely occurring due to HAQER CpG Beacons tending to occur in regions prone to meiotic DNA breaks [46] (15 of 107; p < 10^-8^). To understand if interlocus gBGC is also shaping the HAQER CpG Beacons we investigated if any of these regions have close paralogs within the human genome (Methods). 29 of the 107 HAQERs CpG Beacons have a close paralog in the genome (3-fold enrichment; p < 10^-9^), suggesting that these regions experienced a high rate of weak-to-strong mutations from interlocus gBGC. Taken together, both interallelic and interlocus gBGC are likely shaping the evolution of these regions through both introducing weak-to-strong substitutions and pushing them to higher allele frequencies in the human population.

gBGC has been proposed as one mechanism contributing to regions in the genome undergoing accelerated rates of substitution, and specifically to the origin of human accelerated regions (HARs) [47–49]. If the rapid gain of CpG sites was purely due to gBGC, we might expect a similar increase in GpC sites, which should be created at a rate similar to CpGs when there is a general preference for weak-to-strong substitutions. We observed a ratio similar to this theoretical expectation across phastBias regions not overlapping HAQER CpG Beacons where there were 1.08 CpG gains for every GpC gain. Interestingly, HAQER CpG Beacons showed a specific bias for gaining CpG sites over GpC sites, averaging 1.46 CpG gains for every GpC gain. Because the HAQER CpG beacons were ascertained specifically for a gain of CpGs, we repeated this analysis across thresholds of human-specific GC gain, agnostic to dinucleotide order. A region passed the threshold if the GpC gains or CpG gains passed the threshold, considered separately. This analysis revealed that above a GC dinucleotide threshold of 35 gains per kb, there was a greater than 1.5-fold enrichment of CpG gains compared to GpC gains (Fig. 5f). This is a significant departure from the expected ratio in sites being acted on by gBGC alone, here represented by the phastBias regions. This result is consistent with both gBGC and a selective pressure for gaining CpG sites having shaped the HAQER CpG Beacons, with the strongest examples including HAQER0223 (65 CpG gains vs. 24 GpC gains) and HAQER0020 (97 CpG gains vs. 13 GpC gains) (Fig. S5d). Also consistent with selection, we observed an increased divergence to polymorphism ratio of CpG sites in HAQER CpG Beacons compared to the general set of phastBias regions (Fig. 5g). A mixture of gBGC and selection has been proposed as shaping a subset of HARs [47], and the results of our analyses also point to a mixture of gBGC and a selective pressure for specifically CpG gains, shaping the divergence of HAQER CpG Beacons.

The role of high density CpGs in creating permissive chromatin nominates these variants as increasing the gene regulatory potential of the HAQER CpG Beacons [19]. Indeed, HAQER CpG Beacons were 7-fold enriched among the 50 brain-accessible HAQERs we detected, compared to the set of 1,581 HAQERs (p <2.2x10^-16^) (Fig. 5h). In addition, HAQER CpG Beacons were more likely to be differentially accessible between human and rhesus compared to other brain accessible HAQERs (p=0.00033) (Fig. 5i, Fig. S5f). Finally, HAQER CpG beacons were over 500-fold enriched for overlaps with human-specific H3K4me3 peaks in adult prefrontal cortex from previous studies [38] (p < 10^-37^) (Methods), and significantly overrepresented in human-enriched H3K4me3 peaks compared to rhesus in developing cortex (Fig. 5j). Notably, both HAQER0223 targeting *EMILIN2* and HAQER0074 targeting *SLC35F5* showed a preference for gaining CpG over GpC sites and also showed human-specific gain in regulatory activity in reporter assays during mouse cortical development [8]. These findings support a model in which both gBGC and selection achieve a high density of CpG dinucleotides, which changes the epigenetic state of the chromatin, influencing the expression of nearby genes.

Several of the strongest HAQER-gene linkages we detected reflected this pattern of significant CpG gain in the human lineage. We found that HAQERs 0297 and 0202 regulate transcripts at the *CHL1* locus and that 0261 and 0710 regulate transcripts at the *DPP10* locus, with all of them being HAQER CpG Beacons (Fig. 5k). Furthermore, HAQERs 0297, 0261, and 0710 overlapped human-specific neuronal H3K4me3 histone modifications in the PFC [38,39]. HAQER-gene linkages identified by our study further highlighted that HAQER0261 regulates the *DPP10-AS1* transcript and HAQER0710 distinctly regulates the *DPP10* transcript. Notably, the *DPP10-AS1* transcript is described as a human-specific noncoding gene that reduces the expression of the short isoform of *DPP10* in the prefrontal cortex neuronal layers [38]. We nominate this locus for further study as an example of gBGC, visible to selection, that altered gene regulation in the human cortex.

## Discussion

This study links HAQERs to their target genes in their native genomic context during cortical development and supports gBGC-driven CpG gain and selection as forces forging new gene regulatory functions. While previous studies demonstrated that human-specific sequence changes alter the capacity of HAQERs to drive reporter gene expression during mouse corticogenesis [8], we provide evidence that these fast evolving sequences are necessary to regulate gene expression at their native locus in primary cells from human cortical development. Using an all-in-one CRISPRi system in primary cortical cultures, we epigenetically repressed 50 HAQERs that are accessible during human brain development. We observed resulting down-regulation of nearby candidate *cis-*target genes within a 0.5-Mb window for 26 HAQER perturbations, including coding, antisense and long non-coding transcripts. A subset of HAQER-gene pairs exhibited cell type specificity and/or context dependence under CHIR stimulation, indicating that HAQER activity can be lineage-restricted and/or Wnt pathway responsive. These features are consistent with prior reports that HAQERs are enriched for bivalent chromatin in neurodevelopmental tissues, a state associated with precise spatiotemporal control of transcription and the ability to quickly respond to the cell’s environment, such as developmental signals [8]. Finally, we provide a window into the sequence origins for a subset of HAQERs that were shaped by GC-biased gene conversion, introducing a set of 107 HAQER CpG Beacons enriched for regulatory functions during human cortical development.

Our results align with a model in which DNA sequence changes in noncoding elements alter gene regulatory potential and interact with epigenetic layers to drive transcriptional differences [50]. These changes in genetic and epigenetic layers have been hypothesized to fine tune gene expression changes in developmental time and anatomical location, driving phenotypic divergence [50–54]. CpG accrual can create DNA hypomethylation and subsequent binding of factors associated with active transcription [19,40,41]. Consistent with this model, HAQER CpG Beacons contain dense clusters of human-specific CpG dinucleotides and are enriched for DNA hypomethylation, CFP1-binding, chromatin accessibility, differential accessibility during cortical development, and human-specific H3K4me3 in prefrontal cortex. Notably, several of the strongest HAQER-gene linkages at the *CHL1* and *DPP10* loci overlap with multiple HAQER CpG Beacons and human-specific H3K4me3 marks in prefrontal cortex. HAQERs 0710 and 0261 in the *DPP10* locus pair with distinct transcripts *DPP10* and the human-specific *DPP10-AS1* transcript, respectively, suggesting that HAQER0261 may be rewired to regulate *DPP10-AS1* in humans. The human-specific *DPP10-AS1* transcript is specifically expressed in neuronal layers in the PFC, highlighting species and cell-type specificity in HAQER0261 regulatory function [38]. Similarly, HAQER0223 represents another HAQER CpG Beacon with strong linkage to a downstream gene, *EMILIN2,* that also shows a gain of human-specific regulatory activity in reporter assays [8].

While researchers may have previously viewed forces that increase mutation rates or meiotic drives as mutually exclusive with selection, several observations support the joint action of gBGC and selection in HAQER CpG Beacons. These results include: the directed gain versus loss of CpGs, the preference for CpG over GpC dinucleotide gains, an increased CpG divergence to polymorphism ratio compared to other gBGC loci, the strong overlap with human-specific neuronal H3K4me3 acquisition, the tendency to be accessible in cortical progenitors, and the functional gene-regulatory effects in primary cells. This view of gBGC being compatible with selection is consistent with studies showing that human-accelerated regions are better fit by models that include both gBGC and changes in the selection coefficient [47]. Similarly, we propose that the patterns we observe in HAQER CpG Beacons are most consistent with selective pressure acting on a subset of gBGC driven CpG-gain mutations, resulting in gene regulatory innovations on the human lineage

Beyond the idea that gBGC and selection are able to work in concert, we propose that these two forces may be inclined to jointly forge functional elements in our genomes. Selection can not act on its own, but rather needs mutations to favor. This observation has led to a long-standing interest not only in the "survival of the fittest," but also the "arrival of the fittest" [55] to understand what types of process are likely to introduce advantageous variation. Interlocus gBGC is a source of elevated mutation rates, and more variants being produced in a region may bias that region to be the source of an advantageous variant over a particular length of time. While selection can act on a variant as soon as it has arrived in the population, selection can be inefficient when alleles are at extreme frequencies, such as when a variant first enters the population. Interallelic gBGC may not be an initial source of mutations, but it can contribute to variants arriving at an allele frequency where selection can more efficiently act on them. Interallelic gBGC can also produce novel combinations of alleles when the individual alleles are already present in the population. Previously, we may not have held an *a priori* bias that a cluster of adaptive alleles would heavily favor weak-to-strong mutations, and we may have even viewed such a signal as evidence purely gBGC was occurring at a locus; however, the model that CpG dense regions are primed for proto-enhancer function [19] supports large numbers of weak-to-strong mutations being a favorable path for producing gene regulatory elements. Along with these features of gBGC that appear poised to efficiently forge regulatory regions, we also noticed that, while we did not ascertain regions for functional testing based on CpG gains or weak-to-strong mutations, regions with these features were enriched for accessible DNA, human-specific epigenetic features, and gene regulatory function at the native locus in our CRISPRi assays. Taken together, we entertain the idea that not only is it possible for gBGC and selection to co-occur, but that the gBGC mechanism may be an efficient source of new gene regulatory elements. While our observations focused on the human lineage, this mechanism could potentially be acting more broadly across mammals to regulate chromatin accessibility and gene expression.

Two features of our design strengthen inference about physiologically and clinically relevant causal regulation in human brain development. First, assaying primary human cells preserves the physiological chromatin and transcription-factor landscape that governs cell type specific enhancer function. Second, an all-in-one Perturb-seq technique directly captures guide identity and the transcriptome in a given cell enabling multiplexing perturbations while confidently capturing subtle *cis-*regulatory effects typical of noncoding perturbations. Nevertheless, the activity of sgRNAs can vary depending on their genomic position, [28,56,57], Perturb-seq is more well powered for detecting changes in abundant genes [28,58], and we ascertained candidate HAQERs mainly in radial glia under baseline conditions, indicating that the set of HAQER-gene linkages we identified here is not exhaustive. In addition, although CRISPRi typically exhibits focal specificity,[59] it is possible that targeting may result in additional phenotypes from spread of repressive heterochromatin outside of the HAQER. Future analyses with emerging T2T genomes, comparative multiomics atlases, epigenetic signatures, and additional developmental stages and stimulation contexts will further refine the search for functional HAQERs affecting human brain development and function.

## Conclusions

Together our study informs the identity, extent, and genomic origins of HAQERs influencing gene regulation during cortical development, highlighting an underappreciated role of gBGC as a mechanism supporting the fixation of new gain-of-function functional regulatory elements that may influence human brain evolution. Our findings provide insights into global mechanisms of enhancer evolution and open the door to future prioritization and mechanistic studies of individual candidates, including the clinically relevant *CHL1* and *DPP10* loci, each harboring pairs of HAQER CpG Beacons.

## Acknowledgements

The authors acknowledge and thank Jingwen Ding, Emily Corrigan, Tyler Fair, Matthew Schmitz, Lauretta El Hayek, Reed McMullen, and other Pollen lab and Lowe lab members for valuable comments and bioinformatic resources, Nathan Schaefer for supporting sgRNA and individual assignments, J. Srivastava and K. Pastores for their assistance performing cell sorting at the Gladstone Flow Cytometry Core, supported by NIH S10 RR028962, S. Wang for transferring samples, and A. Tarantal for providing samples. We acknowledge the following funding sources: U01MH114825 and UM1MH130981 (A.A.P.), R01MH134981 (A.A.P.), R35HG011332 (C.B.L.), the Chan Zuckerberg Biohub (A.A.P.), an Ann B. Bussel Research Award from the Ruth K. Broad Foundation (C.B.L.), Schmidt Science Fellows and Jane Coffin Childs fellowships (J.L.W.) and the Duke/UNC ADRC P30AG072958, National Institutes of Health DP2MH122400-01, Schmidt Futures Foundation, Shurl and Kay Curci Foundation Innovative Genomics Institute Award, W.M. Keck Foundation, and William K. Bowes Jr. Foundation. A.A.P. is a New York Stem Cell Foundation Robertson Investigators and a member of the UCSF Kavli Institute for Fundamental Neuroscience.

## Author Information

### Authors and Affiliations

The Eli and Edythe Broad Center of Regeneration Medicine and Stem Cell Research, University of California San Francisco, San Francisco, CA, USA

Yashodara Abeykoon, Enakshi Sinniah, Bryan J. Pavlovic, Jenelle L. Wallace, & Alex A. Pollen

Department of Neurology, University of California San Francisco, San Francisco, CA, USA

Yashodara Abeykoon, Enakshi Sinniah, Bryan J. Pavlovic, Jenelle L. Wallace, & Alex A. Pollen

Weill Institute for Neurosciences, University of California San Francisco, San Francisco, CA, US

Alex A. Pollen

Department of Molecular Genetics and Microbiology, Duke University, Durham, NC 27710, USA

Natalie Dzikowski, Yanting Luo, Seth Weaver, Riley J. Mangan, Craig B. Lowe

Department of Cell Biology, Duke University, Durham, NC 27710, USA

Yanting Luo, Seth Weaver, Craig B. Lowe

University Program in Genetics and Genomics, Duke University, Durham, NC 27710, USA

Natalie Dzikowski, Yanting Luo, Seth Weaver, Craig B. Lowe

Present address: Computer Science and Artificial Intelligence Laboratory, Massachusetts Institute of Technology, Cambridge, MA 02139, USA

Riley J. Mangan

Department of Integrative Biology, University of California, Berkeley, California, USA

Landen Gozashti, Peter H. Sudmant

## Contributions

Y.A. C.B.L., and A.A.P. conceived the project and experimental design. C.B.L. and A.A.P. supervised the study and secured the funding. Y.A. established the all-in-one vector, performed all the experiments, and analyzed the data with the help of R.J.M., Y.L. and E.S. for ATAC-seq, J.L.W for Dreamlet, and S.W. for SCEPTRE. J.L.W., and B.J.P. performed initial testing of the primary culture system. N.D. performed CpG enrichment, gBGC analyses, and DNA methylation assays in neurons and oligodendrocytes. Y.L. performed differential accessibility analyses. S.W. performed paralog identification. E.S. performed H3K4me3 enrichment, ATAC peak characterization, and CUT&Tag analyses. L.G and P.H.S performed DNA methylation in human sperm. J.L.W. performed CrossPeak analyses. Y.A., C.B.L., and A.A.P. prepared the manuscript with input from all authors.

## Ethics Declaration

All studies were approved by UCSF GESCR (Gamete, Embryo, and Stem Cell Research) Committee.

## Materials and Methods

### Data availability

Raw sequencing data and processed data will be made available through dbGaP upon publication under phs004522. ATAC data for Rhesus is available through GEO accession number GSE320143. All genomic coordinates reported in supplementary files correspond to the hg38 reference genome unless noted otherwise. Any additional information and code required to reanalyze the data reported in this paper is available from the lead contact upon request.

### Human and Rhesus Macaque Primary Cell Culture

#### Ethics Statement for human tissue acquisition

Developing cerebral cortex tissue from Gestational Week (GW) 14,16,22, and 23 were collected with patient consent under institutional review board regulations of University of California, San Francisco.

#### Ethics Statement for rhesus macaque tissue acquisition

Developing cerebral cortex tissue from Post Conception Day (PCD) 50,60,80, and 85 were collected under institutional review board regulations of University of California, Davis Primate Center.

#### Generation of 2D cell cultures from primary tissue

The Primate Center at the University of California, Davis, provided specimens of cortical tissue from rhesus macaques. Dissections were performed in PBS under a stereo dissection microscope (Olympus SZ61). Presumptive regions were dissected such as motor versus somatosensory cortex before the appearance of the central sulcus or the anterior ends of the MGE and LGE as reported before [60]. Rhesus PCD50 motor cortex, PCD60 motor and parietal cortex, PCD80 parietal cortex, and PCD85 cortex were used in this experiment. De-identified human tissue samples were collected with previous patient informed consent in strict observance of the legal and institutional ethical regulations as reported previously [61–63]. Collected human and rhesus macaque cortical tissue was cut into approximately 2x2 mm^2^ small pieces using a sharp blade, followed by incubation in Hibernate-E (Thermo A1247601) and CEPT cocktail[64] for 1 hour to increase cell viability Hibernate-E and CEPT were removed and replaced with Bambanker freezing media (NIPPON Genetics, BB02) at a 2:1 ratio with the tissue. 1.5 mL of Bambanker and tissue were transferred to each cryovial, placed in a Coolcell and frozen in -80°C. Cryopreserved vials were then transferred to LN2 for long term storage. Prior to 2D plating, cryopreserved tissue vial was thawed in a 37°C water bath. The tissue was then dissociated following Worthington Papain Dissociation Kit manual (Worthington #LK003150). Dissociated cells were plated on 0.1% PEI (Sigma, P3143), 5 μg/ml Biolamina LN521 (Invitrogen, 23017-015) coated plates and cultured in progenitor enriching media: DMEM/F12 (Corning# 10-92CM or Hyclone SH30261.01) supplemented with BSA, Insulin (Thermo, A1138IJ), Transferrin (Invitria, 777TRF029-10G), Selenium (Sigma, S5261-10G), 1.23 mM ascorbic acid (Fujifilm/Wako, 321-44823), Defined Lipids (Thermo 11905031), Nucleosides (ACGU@20 mM, T@8mM), 20ng/mL FGF2-G3/EGF/BDNF, and 100ug/mL Primocin (Invivogen). This media also contained CEPT cocktail when first plating the dissociated cells to boost cell survival. The media was changed every 2-3 days.

### ATAC sequencing

#### Sample preparation

Age matched human and rhesus macaque tissue was dissociated from frozen tissue, plated at an initial density of 300,000cells/cm^2^, and expanded until fully confluent. At passage 1(P1), all samples were dissociated from 2D and cryopreserved as P1 except the PCD50 rhesus sample which was cryopreserved at passage 3 due to the smaller volume of starting tissue. For ATAC seq, all samples were then thawed in parallel in arrayed format and passed once more to confirm cell viability. On the day of ATAC-seq, cells were dissociated with Papain and 2 technical replicates of 100,000 cells for each biological sample were collected. Cells were processed following the Omni-ATAC protocol [65] for library preparation with minor adjustments. Transposed DNA was amplified using i5 and i7 indexing primers using the following PCR program: 72°C for 5 min; 98°C for 30 s; 10 cycles of 98°C for 10 s, 65°C for 75 s; 4°C hold. We purified the resulting libraries with SPRI beads (active motif protocol) and quantified their concentration using bioanalyzer and qubit. All samples were pooled and submitted for multiplexed sequencing using NovaseqX platform.

#### Bulk ATAC-seq preprocessing and alignment

Raw sequencing reads were adapter-trimmed using Cutadapt v4.4, removing Nextera transposase adapter sequences (CTGTCTCTTATACACATCT), low-quality bases (Phred score <25), and reads shorter than 20 bp. Reference genomes were downloaded from UCSC (hg38 for human, rheMac10 for rhesus macaque) and indexed using Bowtie2 v2.5.1. Trimmed paired-end reads were aligned to their respective species-specific genome indices with Bowtie2 using the --very-sensitive preset, allowing up to 10 multi-mapped locations per read (-k 10) and a maximum fragment length of 2000 bp (-X 2000). Only uniquely mapping reads with MAPQ ≥ 30 were retained using Samtools v1.15.

#### BAM file processing and peak calling

Aligned BAM files were sorted either by genomic coordinate (for IGV visualization) or by read name (for Genrich peak calling). Sorting and indexing were performed with Samtools. To identify regions of accessible chromatin, peaks were called using Genrich v0.6 in ATAC-seq mode (-j -r), excluding mitochondrial reads (-e chrM) and applying a q-value threshold of 0.05. NarrowPeak output files were generated for each sample and species.

#### Quality control and replicate correlation

Correlations between technical and biological replicates were assessed using deepTools v3.5.0. The multiBamSummary function was run in BED-file mode, using species-specific union peak BED files as input. Spearman correlation matrices and heatmaps were generated with plotCorrelation to assess reproducibility.

#### Peak annotation and visualization

For intersection with HAQERs and VISTA elements, human and rhesus macaque Genrich narrowPeak files were intersected with curated BED files of genomic elements using Bedtools v2.30.0[66]. Intersections were performed with a reciprocal overlap threshold of 50% (-F 0.5). Cross-species liftOver was used to map HAQER hg38 coordinates to the rheMac10 genome and then intersected with rhesus ATAC peaks to identify HAQERs overlapping accessible chromatin in rhesus samples. BAM files were uploaded for IGV [67] version 2.16.1 for interactive visualization. ATAC-seq signal tracks for HAQERs and VISTA loci were visualized with SparK[68]. BedGraph tracks were generated from position-sorted BAMs using deepTools bamCoverage. In the SparK plots, the x-axis denotes genomic coordinates and the y-axis represents per-base ATAC-seq coverage (reads per 1-bp bin). Heatmaps visualizing average ATAC-seq read density at TSS promoters were generated using the Enrichedheatmap package [69].

#### Human vs Rhesus accessible peaks

Bulk ATAC-seq peaks (Genrich) from the human and rhesus macaque second trimester developing brain were used to identify homologous open chromatin regions between the 2 species. Human (hg38) peaks were lifted to rhesus (rheMac10) and vice versa (kentUtils: liftOver, gonomics: intervalOverlap) [70] [71]. Native peaks were filtered to retain only those that successfully lifted over, ensuring one-to-one correspondence between native and lifted peaks. Filtered and lifted peaks were concatenated within each assembly and overlapping peaks were merged, producing 2 sets of unified merged peaks, one on the human assembly, and one on the rhesus assembly (bedtools:sort, bedtools:merge) [66]. Replicate-level bulk ATAC-seq BAM read counts were quantified over the merged peaks (DiffBind) and normalized using the median of ratios method (DESeq2). The merged peaks were also intersected with HAQER regions (bedtools:intersect-wao) [66]. The resulting count matrices for the merged peaks with HAQER intersection information were then mapped to unmerged peaks to flatten data, ensuring that each pair of homologous human-rhesus peaks had associated read count and HAQER intersection data. A scatter plot was constructed for the ATAC-seq mean normalized read counts over each pair of homologous human-rhesus peaks. Spearman’s rank correlation coefficient and significance were calculated for the mean normalized read counts in human and rhesus. Overlap based enrichment was performed using a fisher’s exact test to compare the fraction of CpG beacons intersecting brain accessible HAQERs.

#### Human vs Rhesus differentially accessible peaks

Bulk ATAC-seq read counts over human-rhesus paired consensus peaks were used to identify peaks that were differentially accessible between the 2 species (DiffBind, FDR < 0.05). Overlap based enrichment was performed using a fisher’s exact test to compare the fraction of differentially accessible peaks intersecting CpG beacon brain accessible HAQERs.

#### Overlap enrichment analysis

Overlap based enrichment was performed using a fisher’s exact test and odds ratio calculation to compare the fraction of VISTA elements categorized by tissue group (*N* = 2314) intersecting with human and rhesus ATAC peaks generated in this study. All VISTA elements were grouped based on their expression in only brain-related tissues (brain-specific), brain-related and other tissue types (brain-mix) or only other non-brain tissues (other tissue). The significance of the overlap between cortical progenitor accessible HAQERs (N = 50), sgRNA level gene-linked HAQERs (N = 26), element level gene-linked HAQERs (N = 17) and human-specific prefrontal cortex H3K4me3 domains (N = 140) or CpG beacons (N = 20) was determined using the regioneR package [72]. Briefly, we ran ‘permTest’ using 10,000 permutations by randomly resampling from a universe of all HAQERs (N = 1581) as a null distribution to determine the number of overlaps between the sets of interest compared to background.

### All-in-one Perturb-sequencing vector and sgRNA design

#### Plasmid construction and quality control

pLV hU6-sgRNA hUbC-dCas9-KRAB-T2a-GFP was a gift from Charles Gersbach[25] (Addgene plasmid #71237; http://n2t.net/addgene:71237; RRID:Addgene_71237) and was used as the parent vector. Its sgRNA scaffold was modified using Vectorbuilder services to insert capture sequence 1 (cs1) to create the All-in-one Perturb-seq vector to enable single cell capture. The efficiency of the all-in-one vector was measured by targeting B2M surface protein in cultured human primary cells using published B2M sgRNA[27]. Briefly, human cortical progenitor cells were plated in 2D and transduced with 4µL/mL concentrated lentivirus containing the all in one CRISPRi vector and either B2M or non-targeting control (NTC) sgRNA. We selected transduced cells using 5μg/ml Bleomycin sulfate(Cayman chemical company, #13877). The cells had to be passed at least once and replated to allow cell death of untransduced cells in order to select for 99% pure GFP+ cells. After 11-15 days of transduction, cells were dissociated using Papain+ DNAse and the concentration was adjusted to 1-5 million cells/ml. 100 μl of this cell suspension was added to 400 μl of PBS+1%BSA and blocked for 30 minutes on ice. Cells were then stained with 1:200 B2M-APC Antibody (Biolegend, Cat#316311) for 15 minutes at room temperature. Cells were washed 3 times in PBS by centrifuging at 300xg for 5 minutes and resuspended in 200ul of ice cold FACS buffer. Cells were then analyzed by flow cytometry and the resulting data were analyzed by FlowJo. Untransduced cells from the same individual were used for gating. B2M surface protein was visualized through APC channel and GFP from the lentivirus was visualized through GFP 488 channel.

#### Vector Digestion

For sgRNA to be cloned in, the All-in-one vector was first digested with BsmBI-v2 at 55°C for 30 minutes (New England Biolabs, R0739). BsmBI was then heat-inactivated at 80°C for 20 minutes. To avoid any self ligation potential of the vector, an alkaline dephosphorylation step immediately followed. The reaction was placed back on ice and 2µl of quick CIP (New England Biolabs, M0525S) was added and incubated at 37°C for 10 minutes followed by inactivation at 80°C for 2 minutes. The digested and dephosphorylated vector was run on a 1% agarose gel at 120V and was extracted using the Zymo Gel DNA Recovery kit (Zymogen, D4002).

#### High MOI sgRNA library design and synthesis

Candidates were prioritized for the high MOI screen by intersecting human bulk ATACseq peaks with HAQER peaks using hg38 genome. 50 candidates were prioritized which had accessible chromatin in at least one human developmental stage. Up to 20 sgRNA sequences for each candidate was designed by Broad CRISPick tool and CRISPOR for hg38 genome. The sgRNA were loaded into IGV and manually inspected for the region of alignment with accessible chromatin of the regulatory element targeted. sgRNA sequences that aligned nearest to the intersection of HAQER and ATAC peak summits were prioritized. 7 known VISTA elements with telencephalon LacZ expression was also included in the Perturb-seq study and CRISPOR was used to design sgRNAs for the vista elements. From CRISPOR, only the sgRNAs with the best efficiency and specificity were chosen denoted by the green color. HAQER elements with higher repetition had lower amounts of sgRNAs per element. An sgRNA pool of 383 with 3-7 sgRNAs per target and 28 non-targeting sgRNAs[27] was synthesized using Twist Biosciences (Table S3) that followed 5’ to 3’ oligonucleotide order of:

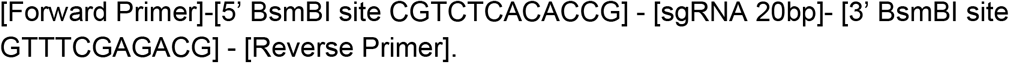

HAQERs were split into either proximal and distal depending on the location of the sequence relative to the nearest TSS. The two categories were synthesized using different forward and reverse primer adaptors and PCR amplified using those respective primers to create separate promoter and enhancer libraries. For sgRNA library amplification, the oligonucleotide template pellet was resuspended in 23 µl of TE buffer (Invitrogen,12090015). 2 µl of template was amplified using 50 μL 2x NEBNext Ultra II Q5 Master Mix (New England Biolabs, M0544S), 5 μL of each 10 μM F and R primer, and 30 μL nuclease-free water. PCR cycling conditions: (1) 98°C for 30 s; (2) 98°C for 10 s; (3) 62°C for 30 s; (4) 72°C for 30 s; (5) go to (2), x 12; (5) 72°C for 2 min. The PCR amplified oligos were cleaned up using the Zymo oligo clean up kit(Zymogen, D4060). Then the oligo pool was digested using Esp3I (New England Biolabs, R0734S) for 1 hour.

#### Low MOI sgRNA library synthesis

29 sgRNAs that were further prioritized by the high MOI screen were synthesized using IDT as separate Forward and Reverse oligonucleotides:

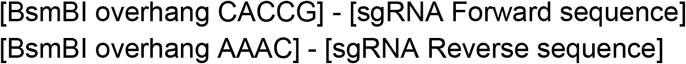

Since the digested vector is dephosphorylated in the previous step, the IDT oligos were phosphorylated to enable successful ligation. Each Forward and Reverse oligo was phosphorylated as a separate 20µL reaction by using 2µL of each 100µM IDT oligo and incubating with 1µL T4 PNK (New England Biolabs, M0201S), 2µL of 10x T4 DNA Ligase buffer, and 15µL sterile water at 37°C for 1 hour followed by inactivation at 65°C for 20 minutes. Phosphorylated oligos were annealed by mixing 5µL of each Forward and Reverse oligos with 90µL of sterile water and incubating in 95°C for 3 minutes followed by room temperature bench top cooling for 1 hour. The annealed oligos were then diluted 1:40 in sterile water.

#### Cloning sgRNA into the All-in-one vector

Both for high and low MOI runs, the sgRNA was cloned into the pre-digested vector as a pool using T4 ligase (New England Biolabs, M0202S) with insert:vector molar ratio of 3:1 The ligated product was transformed using NEB Stable cells(New England Biolabs, C3040H) and plated on LB/Carb(100 μg/mL) plates at 30°C overnight. All the bacterial colonies that grew were scraped into 50 ml of LB Broth with 100 μg/mL Carb and grown overnight. The plasmid was midiprepped using the Zymo midi prep kit (Zymogen, D4200) and submitted to Azenta next generation sequencing(NGS) to confirm the sgRNA library representation. NGS data was analyzed by sgcount pipeline to determine the sgRNAs successfully cloned in.

#### Lentivirus production

Lenti-X 293T cells were plated onto 147cm^2^ dishes coated with Poly-D-lysine (Sigma-Aldrich P7405) and passed at least once in DMEM/F-12 + 10% FBS media. Once the confluence reached ∼80%, media was changed to DMEM/F-12 + 2% FBS. The Lenti-X cells were transfected with the All-in-one plasmid vector together with two viral particle packaging vectors pMD2.G (Addgene, 12259) and psPAX2 (Addgene, 12260) combined at a molar ratio of 2:1:1. The All-in-one vector consisted of Enhancer:Promoter libraries mixed at a ratio of 2:1. Trans-IT transfection reagent (Mirus, MIR 2704) was used to increase transfection efficiency. 12-18 hours post transfection, the media was replaced and ViralBoost (Alstem, VB100) was added to increase viral production. 48 hours post transfection, virus was precipitated using the Alstem system (Alstem, VC100) and resuspended in cold PBS at roughly 1:500 of the original volume. The lentivirus was stored in -80°C and used within one month of production.

#### Primary Cell Transduction

Primary cells were first plated and passed at a density of 150K cells/cm^2^ on PEI + Laminin coated plates. Frozen and thawed lentivirus was mixed with culture media and added on top of cells during feeding. Volume of lentivirus to add was informed by performing a test viral titration step which assessed the percentage of GFP+ cells. Cells were harvested 11 days post transduction for 10x single cell capture. For the high MOI screen, transduction rate was over ∼86% and 10x capture was performed on unsorted cells. For the low MOI screen, transduction rate was ∼10%, and perturbed cells were sorted for GFP+ signal. For both human and rhesus macaque primary cells in our culture, transduction by only adding the lentivirus directly to culture media was important since adding polybrene or viraductin significantly decreased the cell health.

#### Immunocytochemistry

Human and rhesus macaque primary cells were plated in 96-well black, glass bottom plates. Once the cells reached ∼80% confluency, cells were carefully rinsed with PBS once and fixed with 4% PFA for 15 minutes at room temperature. Cells were once again rinsed with PBS twice to remove traces of PFA and blocked with blocking solution PBS + 0.1 % triton-x + 5 % donkey serum for 1 hour at room temperature(RT). Cells were then stained with primary antibodies diluted in blocking solution for 1 hour at RT (TBR1-rabbit-1:300-Abcam-ab31940, GFAP-chicken-1:300-Abcam-ab4674, ki67-mouse-1:600-BD Biosciences-BD55069) Cells were then washed 3 times with PBS, and stained with secondary antibodies for 2 hours. Nuclei were stained with Hoechst (1:10,000, ThermoFisher, 62249). Secondary antibodies were washed off 3 times with PBS and cells were visualized through the EVOS M7000 microscope. Fluorescent channels on microscope -GFAP-488, MKi67-647, TBR1-549.

### Single Cell RNA sequencing

#### High and low MOI 10x sequencing sample preparation

Human and rhesus macaque primary cells were dissociated from 2D culture using Papain+DNAse. Cells were resuspended in PBS+0.1% BSA and either used directly for 10x capture during high MOI assay or sorted for GFP+ cells during the low MOI assays. When sorting was necessary, cells were kept on ice in PBS+1% BSA. The sorter was set to 4°C and the 130 micron large nozzle size was chosen. For 10x capture, cells from all samples were pooled and ∼62K cells/lane were loaded in the high MOI run, or ∼115K cells/lane for the low MOI validation and stimulus response runs. For cell capture and processing, 10x Genomics Chromium X controller and version 3.1 high-throughput RNA kits were used. Gene expression and sgRNA libraries were generated using the manufacturer-provided protocol (CG000421 Rev D).

#### Single cell sequencing and reference genome alignment

Sequencing was performed at the UCSF CAT, supported by UCSF PBBR, RRP IMIA, and NIH 1S10OD028511-01 grants. Cells were sequenced using NovaseqX in either 10B 100 cycles for high MOI run or 25B 300 cycles for low MOI and stimulus response runs. High and low MOI runs each had an approximate sequencing depth of 50,000 and 30,000 reads/cell respectively. sgRNA reads for high vs low MOI runs roughly were around 5000 and 9000 respectively. The resulting sequencing FASTQ output files from high MOI run were aligned to the hg38 genome using Cellranger-7.0.1. Low MOI run was aligned to a chimeric hg38/rheMac10 genome using Cellranger-7.2.0. sgRNAs in each library were aligned using feature barcode references.

#### Demultiplexing of individuals and species

For high MOI run, pooled human samples were demultiplexed using Cellbouncer[73] demux_vcf pipeline on position sorted BAM files output from Cellranger. VCF files needed as input were generated for human samples by SNP genotyping genomic DNA extracted from respective individual samples. Cells called as doublets during individual demultiplexing were removed from further analyses in high MOI data. For low MOI runs, species calls assigned by Cellranger for each cell after aligning to the chimeric hg38/rheMac10 genome were used to separate human from macaque reads. Species doublets were removed from further analyses.

#### sgRNA assignment

For low MOI runs, sgRNAs were assigned to each cell using Cellbouncer demux_tags pipeline. Cellranger output filtered feature barcode matrix was first converted from h5 to mex format using the cellbouncer/utils/h5tomex.py argument. sgRNAs were then assigned to cells using the cell barcodes and features from Cellranger as input together with Cellbouncer argument --feature_type "CRISPR Guide Capture". For low MOI runs, cells were further filtered to have only 1 sgRNA/cell for downstream differential gene expression analysis through Dreamlet.

#### Single cell data preprocessing

We used Scanpy Anndata object structures to preprocess and normalize gene expression data. The adata from different batches in the same run were concatenated. Cells with more than 15% mitochondrial and 40% ribosomal reads were removed. Genes detected in less than 10 cells were removed. Read counts were then normalized, log transformed, and scaled with a maximum value of 10. Principal component analysis (PCA) was then performed on the scaled matrix, and the top 100 principal components were retained. A k-nearest-neighbor graph was constructed from the PCA space using Euclidean distances with k=10 neighbors and the first 40 PCs as input. For the high MOI dataset, batch correction was performed using Harmony integration for batch and individual. Cells were clustered under Leiden resolution of 0.25 using Harmony corrected top 20 PCs and 10 nearest neighbors. For the low MOI datasets, only the genes common to both human and macaque were kept in the Adata object. Integration was performed on species and batch using top 5000 genes in scANVI [74].

#### Single cell data comparison to public datasets and cell type annotation

A publicly available developing human primary cortex multiomic dataset from Wang et.al[61] was used for reference mapping of the high and low MOI datasets separately. Reference data was subsetted to have an equal number of 800 cells representing each cell ‘type’ prior to loading it as the reference in scANVI. The reference model was built with scvi-tools using top 2000 variable genes. The query was either the high MOI dataset or the integrated low MOI dataset. Cell type annotation on Leiden clusters was then performed based on marker expression as well as predictions from scANVI[74].

### SCEPTRE High MOI Differential Gene Expression Analysis

#### Input Variables

To identify gene regulatory effects of enhancer perturbations in a high multiplicity-of-infection (MOI) pooled CRISPRi screen, we applied the SCEPTRE statistical framework using the sceptre R package. Our analysis leveraged an annotated single-cell dataset containing 217,754 cells, of which 67,236 passed stringent cell-level quality control (QC) filters. The screen employed 355 targeting single guide RNAs (gRNAs) covering 50 HAQER and 7 Vista telencephalon enhancer candidate regulatory elements, alongside 28 non-targeting control (NTC) gRNAs. The gRNA-target input table for high MOI SCEPTRE analysis is Table S4.

#### Data Preprocessing and Assignment

Gene expression responses were aggregated at the single-cell level, and both gene and gRNA matrices were jointly subjected to quality control. gRNA-to-cell assignment was performed using a Poisson-Gaussian mixture model, accounting for UMI counts and dropout noise. Assignment covariates included log-transformed gRNA and response detection rates, batch identity, and latent factors inferred by principal component analysis (PC_1 to PC_5).

#### Discovery Analysis Parameters

A total of 1144 candidate gRNA-gene pairs were analyzed, of which 348 passed pairwise QC. For each element-gene pair, we required ≥100 treatment cells (cells with the targeting-gRNA) and ≥100 control cells (cells without that specific targeting-gRNA) with nonzero expression of the response gene. The discovery test used left-sided permutation resampling and a Bonferroni integration strategy to control type I error under high MOI. Multiple hypothesis correction was performed using the Benjamini-Hochberg (BH) procedure at FDR 10%.

Model covariates in the response model included:

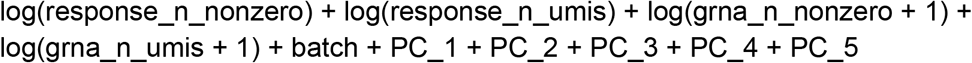

No approximation was used for the permutation resampling. No positive control pairs were defined for this analysis.Out of 348 valid discovery tests, 23 gRNA–gene pairs were identified as significant at FDR ≤ 0.1. Negative control pairs showed no inflation: 0 of 348 NTC pairs were called significant, and the mean log₂FC among NTCs was 0.0062. This indicates proper calibration and specificity of the inference under the high-MOI design.

#### sgRNA level analysis

For sgRNA-level analysis, each guide was tested individually using a singleton integration mode. Here, we applied a more permissive QC requirement of ≥10 treatment and ≥10 control cells with nonzero expression. In total, 3,439 gRNA–gene pairs passed QC. Discovery testing again used permutation resampling with BH correction at FDR 10%. This analysis identified 100 significant gRNA-gene pairs out of 3,439 (5/3,439 NTC pairs significant; mean log₂FC = 0.0093).

### SCEPTRE Low-MOI Analyses (Validation, Baseline, CHIR)

#### Input variables

We analyzed three low multiplicity-of-infection (MOI) pooled CRISPR datasets for Validation, Baseline, and CHIR. Human data was filtered first from integrated single cell objects and used as input for SCEPTRE. Each screen included 25 targeting gRNAs (21 targets) and 4 non-targeting control (NTC) gRNAs. Treatment/control cells were set to 7/7 for validation, and 10/10 for CHIR and Baseline. Four per-cell covariates were available: grna_n_nonzero, grna_n_umis, response_n_nonzero, and response_n_umis. The gRNA-target input table for low MOI SCEPTRE analysis is Table S7.

#### Data preprocessing and gRNA assignment

Gene-level responses and gRNA counts were jointly quality-controlled at the cell level. gRNA-to-cell assignment used the mixture method with the assignment formula:

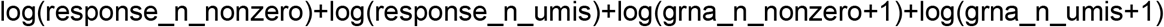

#### Discovery analysis

Discovery tests were left-sided, comparing targeting to NTC cells, with permutation resampling (no approximation). The response model adjusted for:

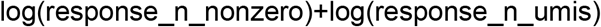

Multiple testing correction was calculated by the Benjamini–Hochberg method (FDR 10%). The integration strategy was singleton.

#### Cell type–specific differential gene expression with Dreamlet

Adata objects were converted to R objects using Zellekonverter. Cell type–resolved differential expression (DGE) was performed with dreamlet v1.3.1 on pseudobulk expression profiles derived from annotated single cells. For preprocessing, raw counts were aggregated to pseudobulk samples within each cell type and processed with processAssay() using min.count = 5, min.cells = 5, and min.prop = 0.20; library-size normalization used the RLE method (norm.method = "RLE"). Only genes meeting these thresholds within a given cell type were retained for testing.

Pseudobulk groupings were set to match the scientific question. For the low-MOI validation analysis, cells were grouped by species × sgRNA to create the species_sgRNA factor. For the CHIR vs Baseline analysis, cells were grouped by species × sgRNA × condition (condition being CHIR or Baseline) to create the species_sgRNA_condition factor. These factors defined the design levels used for inference.

Per cell type and gene, dreamlet fit linear mixed models with a no-intercept parameterization and batch as a random effect:

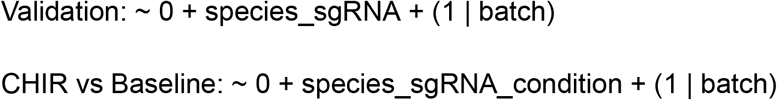

Contrasts compared targeting sgRNA levels to the corresponding non-targeting control within the same condition (species for Validation; species×condition for CHIR vs Baseline). Dreamlet reported effect sizes (log₂FC) and p-values). Unadjusted p values were used for noting significance in the bar plots since each hypothesis for HAQER-gene linkage was formed prior to testing with Dreamlet using BH adjusted analysis from SCEPTRE.

#### HAQER human-specific signatures from previous studies

Human-specific epigenetic signatures were obtained from:

- H3K4me3 PFC tissue-specific and NeuN+ cell-type specific-Shulha et al [38] -hg19 coordinates
- CpG Beacons-Bell et al [39,42] -hg19 coordinates
- Biased Gene Conversions (gBGC)-Dreszer et al, Capra et al [44,45] -hg18 coordinates

Coordinates from previous human genome (hg) versions were lifted over to the hg38 genome for comparisons with HAQER intersections in the main text. Supplementary table S12 reporting the intersections with HAQERs for extreme CpG Beacons clusters from Bell 2012 contain the original hg19 coordinates, and S14 with gBGC contain the original hg18 coordinates reported the respective original studies.

#### Conservation Analysis for PhyloP Panels

To evaluate selective constraint across sets of genomic regions, we accessed two genome-wide, base pair-resolution sequence conservation datasets in UCSC BigWig format representing phyloP scores for each base, quantifying the deviation of observed substitution rates from neutral expectations[75]. These include “Primate PhyloP” scores, estimated from a multiple alignment of 239 diverse primate genomes, and “Mammal PhyloP” scores, estimated from a 447-way alignment of diverse mammals[76,77].

We used the *bigWigAverageOverBed* tool[78] to calculate the mean primate and mammal phyloP scores for each genomic element, and compared the resulting distribution of mean phyloP scores across sets of genomic regions. These include HAQERs, a subset of HAQERs with accessible chromatin in brain, random intergenic regions (excluding ENCODE cCREs, from Mangan et al[8]), ultraconserved elements[77], and human accelerated regions (HARs) [79]. Significant differences in mean conservation is reported as Bonferroni-corrected p values from pairwise t tests.

### CpG Analyses

#### Identification of HAQER CpG Beacons

For each HAQER, we calculated the number of “gained”, “lost” and “maintained” CpG sites in the hg38 reference genome relative to the inferred human-chimpanzee ancestor genome (gonomics: countPairOfBases -compare C G) [71]. A CpG site was considered “gained” if it was present in hg38, but not the human-chimpanzee ancestral genome, “lost” if it was not present in hg38, but present in the human-chimpanzee ancestral genome, and “maintained” if it was present in both genomes. All CpG site changes arising from substitutions, insertions, and deletions were included. GpC site divergence was calculated for every HAQER using the same methods (gonomics:countPairOfBases -compare G C) [71] for downstream analyses. We normalized counts of gained CpG sites to the length of each HAQER to obtain a “CpG gain density,” and HAQERs with a CpG gain density >= 0.017 were called as HAQER CpG beacons (corresponding to the 17/kb threshold for CpG beacon clusters established in Bell et al., 2014 [39]).

To assign significance to HAQER CpG beacons, we first calculated a “genome-wide”, per-base rate of CpG gains in the hg38 genome relative to the inferred human-chimpanzee ancestor genome. We calculated the number of gained CpGs in hg38 (excluding centromeric regions, which exhibited poor alignment) and divided it by the total number of non-gapped, non-centromeric bases in hg38. This value served as the expected probability of CpG gains *p*. We then used the cumulative binomial distribution to calculate the probability of observing *k* or more CpG gains in each HAQER given the total number of bases in each HAQER *n* and the genome-wide probability *p* (R: pbinom). We adjusted p-values for each HAQER by multiplying by the number of 500-bp windows in the genome, as this window size was used to identify HAQERs in the original genome-wide screen (Mangan et al., 2022 [8]). HAQERs with an adjusted p-value < 0.01 were considered to have a statistically significant number of CpG gains relative to the genome-wide background.

To assess whether HAQER CpG beacons are enriched in telomeric/sub-telomeric regions of the genome, we calculated the proportion of CpG beacon and non-beacon HAQERs overlapping the first and last 5MB of each chromosome (bedtools intersect -wa) [66], and evaluated enrichment with Fisher’s exact test. To assess the preference for CpG gains over losses in HAQER CpG beacon clusters, we compared the gained/lost CpG ratios for HAQER CpG beacon clusters (n = 107) and all other HAQERs (n = 1383) in the set with the Wilcoxon rank-sum (Mann-Whitney U) test. HAQERs with 0 gained and lost CpGs were excluded from the analysis.

#### Analysis of CpG site polymorphism in HAQERs

To calculate the number of CpG-related polymorphisms in HAQERs, we first obtained phased GRCh38 1000 Genomes variant calls from 3,202 human samples [80] on chromosomes 1-22 and X, available at the following link:

https://ftp.1000genomes.ebi.ac.uk/vol1/ftp/data_collections/1000G_2504_high_coverage/working/20220422_3202_phased_SNV_INDEL_SV/. We filtered the variant data to retain only bi-allelic substitution variants (gonomics: vcfFilter -biAllelicOnly -substitutionsOnly) [71]. Using these variants, we reconstructed an “alternate” hg38 sequence where the reference allele at each variant position was replaced with the alternate allele (gonomics: vcfToFa) [71]. This alternate hg38 sequence was appended to the reference hg38 sequence to find CpG site differences due to polymorphism. We found the coordinates of all CpG site differences between the reference and alternate hg38 sequences genome-wide (gonomics:locateCG -compare) [71], and these were called “polymorphic” CpGs. We repeated this analysis on the aligned hg38 and human-chimpanzee ancestor genomes to obtain the coordinates of all CpG gains/losses in hg38 due to substitutions only. These were called “divergent” CpGs in our analysis. We filtered the polymorphic and divergent CpG sites to retain only those within HAQERs (bedtools intersect -wa -wb) [66], and obtained counts of polymorphic and divergent CpG sites in each HAQER. HAQERs on the Y chromosome were excluded.

#### phastBias gBGC tract divergence & polymorphism analysis

We calculated the overlap enrichment of HAQER CpG Beacons in phastBias gBGC tracts from Capra et al., 2013 [45] using gonomics:overlapEnrichment [71]. After observing a significant enrichment of HAQER CpG beacons in these regions of gBGC, we evaluated several sequence features of phastBias tracts. We calculated CpG and GpC site changes in the human lineage in phastBias tracts following the same methods for HAQERs (gonomics:countPairOfBases -compare) [71]. To assess the preference for CpG gains over GpC gains in HAQER CpG beacons compared to phastBias tracts, we summed the counts of gained CpG and GpC sites within HAQER CpG Beacons (n = 107) and within phastBias tracts that did not overlap HAQER CpG Beacons (n = 9737). For each set, we computed the ratio of summed CpG gains:GpC gains. To address ascertainment bias for HAQER CpG beacons, we then selected HAQERs by varying thresholds of CpG gain densities (>17/kb, >20/kb, >25/kb, >30/kb, >35/kb, >40/kb), or GpC gain densities passing those same thresholds, selecting for regions with a method agnostic to the dinucleotide order. For each threshold, we created a pooled subset of HAQERs passing the threshold for CpG gains and HAQERs passing it for GpC gains, and performed the same calculation of summed CpG:GpC gain ratio for each subset. HAQERs passing the threshold for both CpG gains and GpC gains were counted once when pooling. Fisher’s exact test was used to assess significance of each category of HAQERs relative to phastBias gBGC tracts.

To analyze the level of CpG divergence vs. polymorphism in phastBias tracts, we first filtered the genome-wide sets of “polymorphic” and “divergent” CpG sites in hg38 due to substitutions to retain only those within phastBias tracts (bedtools intersect -wa -wb) [66] and counted the number of “polymorphic” and “divergent” CpG sites in each phastBias region. To evaluate whether HAQER CpG beacons exhibit a higher ratio of divergent to polymorphic CpG sites than phastBias tracts, we compared the divergent/polymorphic CpG ratios for HAQER CpG beacons (n = 107) and non-overlapping phastBias tracts (n = 9297) using the Wilcoxon rank-sum (Mann-Whitney U) test. HAQERs and phastBias tracts with 0 divergent CpG sites and 0 polymorphic CpG sites were excluded from the analysis.

#### Identification of paralogs

First, a self-alignment of the human T2T genome was performed with lastz [81] using the human.chimp.v2 scoring matrix [82], and alignment parameters for closely related species (O=600 E=150 T=2 M=254 K=4500 L=4500 Y=15000 C=0). We then lifted the HAQER CpG Beacons from the hg38 assembly to the human T2T assembly using kentUtils:liftOver [70]. Next, we used gonomics:overlapEnrichment [71] with the normalApproximate setting to calculate the enrichment of human T2T-lifted HAQER CpG Beacons in regions that had a significant self-alignment in the human T2T assembly.

#### Human-specific prefrontal cortex H3K4me3 enrichment

HAQER CpG Beacon coordinates were intersected (bedtools intersect) with an independently published set of human-specific H3K4me3 peaks in adult prefrontal cortex [38], lifted over from hg19 to hg38 (406 of 410 peaks successfully lifted). The observed number of overlapping peaks was compared to the number expected under random genomic placement, yielding 16 observed overlaps against an expected 0.03 (506-fold enrichment; p < 10⁻³⁷).

#### Comparing methylation across regions in sperm, neurons, and oligodendrocytes

We analyzed methylation profiles from bulk sperm PacBio HiFi sequencing of three individuals. Sequencing data for these sperm samples are not permitted to be released for reasons of identifiability. Those wishing to request access to these data may contact Peter Sudmant, who will respond within two weeks. For each sample, we first aligned reads to the human T2T-CHM13v2.0 using pbmm2 (version 1.17.0; https://github.com/PacificBiosciences/pbmm2) with flag --preset CCS and sorted aligned bam files with samtools sort (version 1.17) (https://doi.org/10.1093/bioinformatics/btp352). Then, we inferred site methylation probabilities from mapped HiFi reads in each sample using aligned_bam_to_cpg_scores (version 3.0.0) with default settings (https://github.com/PacificBiosciences/pb-CpG-tools). We used bedtools (version 2.31.0) (https://doi.org/10.1093/bioinformatics/btq033) to intersect per-site discretized modification scores with regions of interest as well as background regions.

We then analyzed methylation profiles from adult human (age 60 and 80 years) neurons and oligodendrocytes from post-mortem dorsolateral prefrontal cortex (DLPFC) samples. Nuclei were isolated from 500mg fresh frozen brain tissue according to the protocol described in Shireby et. al, 2022 [<u>cit]</u>. Neuronal and oligodendrocyte nuclei were stained with 1:1000 Anti-NeuN antibody (Millipore, Cat#:MAB377X) and 1:150 Anti-SOX10 antibody (R&D Systems, Cat#:NL2864R), respectively, and sorted via fluorescence-activated nuclei sorting on the BD FACSymphony™ S6 Cell Sorter. High-molecular weight DNA was extracted from each nuclei population following the protocol and reagents provided in the CUTANA™ Fiber-seq Kit. PacBio whole-genome libraries were prepared following the low-mass protocol for the SMRTbell prep kit 3.0 and subsequently sequenced on the PacBio Revio System. Reads were aligned to the T2T-CHM13v2.0 human reference genome and sorted using pbmm2 (version 26.1.99; https://github.com/PacificBiosciences/pbmm2). CpG site methylation probabilities were inferred using pb-CpG-tools aligned_bam_to_cpg_scores (version 3.0.0) with default settings.

Coordinates of HAQERs (CpG beacons and non-beacons) and phastBias tracts were lifted over to the T2T-CHM13v2.0 genome using kentUtils:liftOver. CpG sites with inferred methylation probabilities were intersected with HAQER CpG beacons, non-beacon HAQERs, and phastBias tracts using bedtools (version 2.30.0). A “background” dataset of CpG sites was generated by running gonomics:simulateBed on the T2T-lifted HAQER CpG beacons (n=112) for 1000 iterations to generate a BED file of random genomic regions with the same average length as HAQER CpG beacons. We inferred the methylation scores of CpG sites within these background regions using the methods described previously. The background set for every sample was slightly different, but all averaged around 1 million (Individual #1 (60y) Neurons: 1012236, Oligodendrocytes: 1015243. Individual #2 (84y) Neurons: 981702, Oligodendrocytes: 964866.) We compared the methylation scores for CpGs within HAQER CpG beacons, non-beacon HAQERs, phastBias tracts, and the background set for each sample using the Wilcoxon rank-sum (Mann-Whitney U) test.

#### HAQER CpG Beacon enrichment in CFP1 binding sites

Using publicly available CFP1 ChIPseq data from human erythroid cells [83], we found that HAQER CpG beacons show greater enrichment of CFP1 binding sites than non-beacon HAQERs and regions of GC-biased gene conversion (phastBias tracts). We aligned the input control DNA and the DNA from cells treated with an anti-CFP1 antibody to the hg38 reference genome using bowtie2 and called CFP1 binding peaks using macs3 (q<0.01). We then ran gonomics:overlapEnrichments to calculate the enrichment value of HAQER CpG beacons for CFP1 binding peaks, and repeated this analysis for non-beacon HAQERs and phastBias tracts.

### H3K4me3 CUT&Tag sequencing

#### Sample preparation & CUT&Tag assay

Bulk CUT&Tag [reff; Kaya-Okur et al, PMID: 32913232] targeting H3K4me3 on human (GW23) and rhesus macaque (PCD85) primary cortical cells (one biological replicate with 2x technical replicates each). Primary cortical samples were cultured and cryopreserved as described in section ‘Generation of 2D cell cultures from primary tissue’. Briefly, cryopreserved vials were thawed in 37°C water bath and cells were centrifuged at 300xg for 3 minutes and resuspended in 1mL of ice-cold PBS, for a total of two washes. Cells were counted and 2 technical replicates of ∼400,000 cells per biological sample were aliquoted. Samples were processed following the CUT&Tag-IT kit manufacturer’s protocol (Active Motif, 53160) with minor changes. Following concanavalin A bead binding, nuclei were incubated with ∼2ug primary anti-H3K4me3 antibody per reaction (1:25, Abcam, ab8580) overnight at 4°C on an orbital shaker. The next day, nuclei were washed and incubated with guinea pig anti-rabbit secondary antibody (1:100, Active Motif, 53160) for 1 hour at room temperature on an orbital shaker. After additional washes, the nuclei were incubated with CUT&Tag-IT-assembled pA-Tn5 Transposomes for 1 hour at room temperature. Tagmentation was performed by incubating samples in tagmentation buffer for 1 hour at 37°C. To stop tagmentation and release tagmented DNA, the samples were treated with EDTA, SDS and proteinase K at 55°C for 1 hour. DNA was purified using columns provided in the kit. Library amplification was performed using indexed primers based on the manufacturer’s guidelines and libraries were size-selected using SPRIselect beads (Beckman Coulter). The final CUT&Tag libraries were assessed for fragment size distribution using Agilent Tapestation and sequenced on an Illumina NovaSeq X plus platform with paired-end reads (R1: 151 bp; BC1: 12bp; BC2: 24 bp; R2: 151 bp).

#### Bulk CUT&Tag data preprocessing and alignment

Raw sequencing read quality was assessed via FastQC v0.12.1. Paired-end reads were adapter-trimmed using Cutadapt, removing Nextera transposase adapter sequences, followed by fastp to remove poly-G tails. Reference genomes (hg38 for human, rheMac10 for rhesus macaque; CAT[reff; Fiddes et al. 2018, PMID: 29884752]-processed, with nuclear mitochondrial insertions masked) were indexed using Bowtie2 v2.5.4. Trimmed reads were aligned to their respective species-specific genome indices with Bowtie2 using the --very-sensitive preset in end-to-end mode, discarding unpaired and discordant alignments (--no-mixed --no-discordant --no-unal) and restricting fragment length to 10–700 bp (-I 10 -X 700). Aligned reads were sorted and indexed with Samtools v1.21, and filtered to a minimum mapping quality of 2. PCR duplicates were retained, consistent with standard CUT&Tag practice, as apparent duplicate fragments in high-quality libraries predominantly represent true independent tagmentation events rather than PCR artifacts.

#### BAM file processing and peak calling

Filtered BAMs were name-sorted and converted to BEDPE using Bedtools v2.31.1, restricted to same-chromosome fragment pairs <1 kb, and reduced to fragment coordinates. Fragment coordinates were converted to genome-wide bedgraph coverage (bedtools genomecov -bg) per replicate and per replicate-merged BAM (samtools merge). Peaks were called from bedgraph tracks using SEACR v1.3 in normalized, stringent mode, run separately on each technical replicate and on the replicate-merged track per species. Merged-replicate peak calls were used as the input peak set for downstream cross-species consensus peak generation.

#### Quality control and replicate correlation

Fragment length distributions were extracted from properly paired, mapped reads to assess nucleosomal periodicity. Peak number, peak width, and reproducibility between technical replicates (fraction of overlapping peaks via GenomicRanges) were calculated for each sample. Fraction of reads in peaks (FRiP) was computed using chromVAR::getCounts on each replicate BAM and its matched peak set. Genome-wide reproducibility was further assessed by binning the genome into 500-bp windows, counting fragment midpoints per bin, and calculating Pearson correlations of log2-transformed bin counts between samples, visualized as a hierarchically clustered correlation heatmap.

#### Peak annotation and visualization

Genome-wide signal tracks were generated from replicate-merged BAMs using deepTools bamCoverage (RPKM-or CPM-normalized, 10-bp bins). H3K4me3 enrichment at TSS and at peak summits was visualized as heatmaps and aggregate profiles using deepTools computeMatrix (reference-point mode) and plotHeatmap/plotProfile, using species-matched TSS coordinates from CAT-lifted[reff; Fiddes et al. 2018, PMID: 29884752] GTF annotations. TSS enrichment scores were calculated per species by normalizing the aggregate TSS profile to mean flanking signal (100 bp at each end of a ±3-kb window), following ENCODE conventions.

#### Human vs Rhesus consensus H3K4me3 peaks

Orthologous consensus H3K4me3 peaks between human and rhesus macaque, were identified using CrossPeak [reff; https://github.com/jenellewallace/CrossPeak]. Briefly, a Progressive Cactus whole-genome alignment (HAL file) spanning eight primates was used, selecting the "primate" node as the human–rhesus coordinate bridge. Using halLiftover, summit-centered, fixed-width H3K4me3 peaks of 501 bps were reciprocally projected between species. Peaks overlapping species-specific blacklist regions or non-standard chromosomes were excluded. Consensus peaks were resolved through a two-round liftover-and-reconciliation procedure (11-bp and 51-bp summit windows for rounds 1 and 2; indel tolerances of 5, 50, and 450 bp), producing one-to-one orthologous consensus H3K4me3 peaks and species-specific H3K4me3 peak sets. Read counts over the merged-replicate consensus peak set were quantified from each technical replicate BAM individually (unmerged) using deepTools multiBamSummary (BED-file mode).

#### Human vs Rhesus differentially enriched peaks

Differential H3K4me3 enrichment over consensus peaks was tested using DESeq2 on the replicate-level count matrix (peaks filtered to summed count ≥10 across samples, species as sole design variable). Peaks with adjusted p < 0.05 (Benjamini–Hochberg) were called differentially enriched; log2 fold changes were shrunk using apeglm. Differential peaks were exported in both human and rhesus genomic coordinates. As only one donor was profiled per species, technical replicates were used in place of biological replicates for dispersion estimation in DESeq2, and species is therefore confounded with individual. Consequently, reported significance estimates should be interpreted with this limitation in mind.

#### Overlap enrichment analysis

Overlap-based enrichment was performed using Fisher’s exact test to compare the fraction of human-specific H3K4me3 peaks (differentially human-enriched consensus peaks plus human-only non-orthologous peaks) intersecting HAQER/CpG beacon coordinates (bedtools intersect), against all remaining consensus peaks as background. Enrichment was corroborated with a permutation test (1,000 permutations of size-matched consensus peak sets). A parallel Fisher’s exact test compared cortex-accessible versus non-accessible HAQERs for overlap with an independently published human-specific prefrontal cortex H3K4me3 peak set [38].

## Supplementary Figures

**Fig S1:**
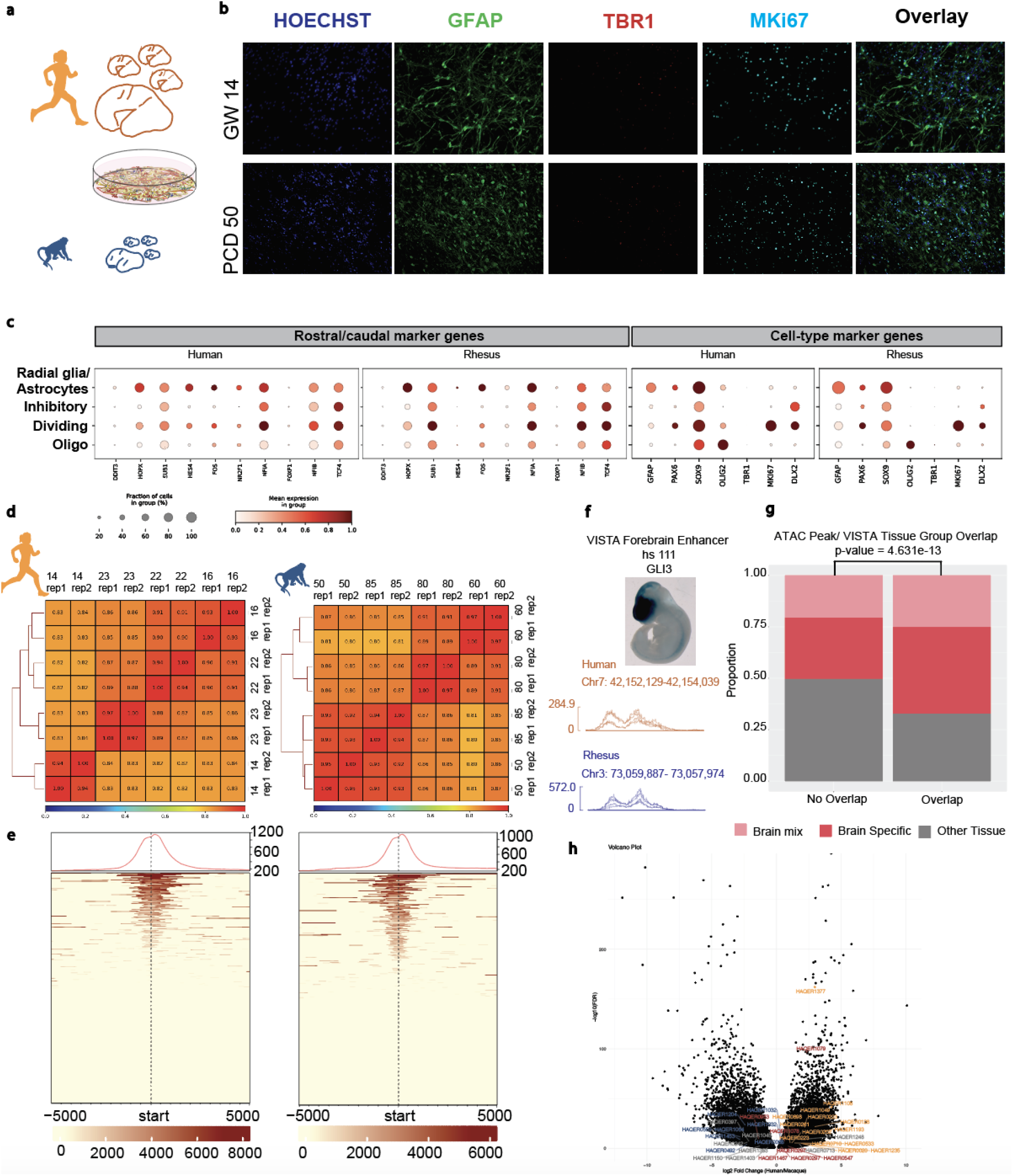
Supplemental analyses of sample information and bulk ATAC sequencing for human and rhesus. a. 4 human and 4 rhesus samples were used throughout the study. Primary cortical tissue was dissociated and plated in 2D. b. Representative samples from GW14 human and PCD50 rhesus immunocytochemically labeled with antibodies against GFAP, TBR1, and MKI67. Nuclei stained in Hoechst. Note that most cells are progenitors labeled with GFAP with minimal TBR1 stained cells. GW= Gestational Week, PCD= Post conception day. c. To assess the expression of marker genes for cerebral cortex areas, scRNA-seq data from the same exact samples used in ATAC-seq and RNA-seq were used which provide molecular subtype resolution on the samples used for CRISPRi experiments. Both human and rhesus samples contain a mixture of cortical areas with similar expression patterns of rostral and caudal markers [62]. Both human and rhesus samples contain cells expressing progenitor markers GFAP, and SOX9. Oligodendrocyte labeled cell clusters express OLIG2, dividing cells express MKI67. Minimal level of TBR1 is present in scRNA-seq data matching the immunocytochemistry results. d. Heatmaps showing correlation between 4 biological replicates for each species. Human samples are GW14, 16, 22, and 23. Rhesus samples are PCD50, 60, 80, and 85. Rep1 and rep2 refer to 2 technical replicates per each sample used for ATAC seq. e. Average ATAC-seq read density at TSS promoters of ENSEMBL genes of human and rhesus developing brain samples. f. Example of element hs111 with lacZ staining in blue in the developing mouse forebrain from VISTA Enhancer Browser (top). Accessibility of hs111 in human and rhesus developing cortex samples via bulk ATAC sequencing visualized as traces for all individuals per species (bottom). Note that hs111 shows significant chromatin accessibility in both human and rhesus developing brain samples. g. Overlap fraction of VISTA elements categorized by tissue group (brain-specific, brain-mix or other tissue) overlapping with accessible chromatin peaks from developing brain samples in this study. h. Volcano plot of differential chromatin accessibility between human and rhesus macaque. Each dot represents an orthologous accessible region identified by the CrossPeak [84] pipeline, which harmonizes ATAC-seq peaks across species. The X-axis shows the log2 fold change of accessibility in Human/Macaque, and the Y-axis shows the –log10(FDR) from differential accessibility testing. Peaks overlapping the HAQERs which are detected to be accessible by the Genrich pipeline, labeled for human-only accessibility in orange, rhesus-only in blue, and shared in red.

**Fig S2:**
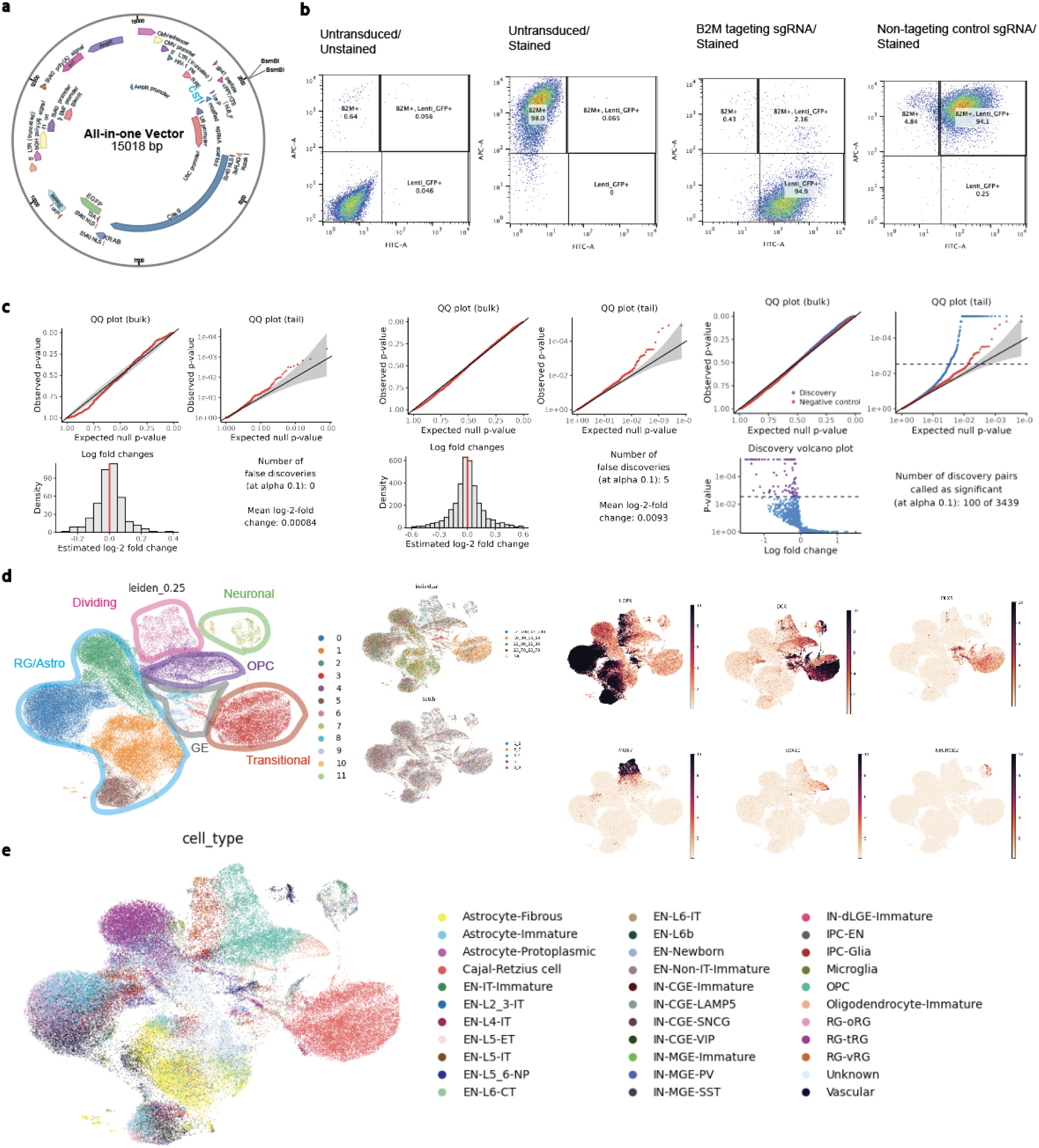
All-in-one Perturb-seq vector establishment and supplemental details for the high MOI screen. a. All-in-one Perturb-seq vector map. Each sgRNA is cloned into the BsmBI restriction site. Expression of EGFP cassette allows visualization and sorting of transduced cells. b. Quality control assay for determining the efficiency of the all-in-one Perturb-seq vector. Cultured human primary cortex cells from GW17 cells were transduced with sgRNA targeting surface protein B2M or non-targeting control. Distribution of cells display APC signal in the Y axis from anti-B2M staining, and FITC signal in the X axis, from EGFP expression by the vector. Left to right: Cells untransduced and not labeled with anti-B2M. Cells untransduced, but labeled for surface marker B2M. Cells transduced with B2M targeting sgRNA and labeled with anti-B2M. Cells transduced with NTC sgRNA and labeled with anti-B2M. c. Left to right: Scatter plots generated by SCEPTRE during calibration step showing zero false discoveries at 10% BH FDR for element level analysis and 5 false discoveries at sgRNA level analysis. Histograms show estimated log_2_FC for paired genes during calibration step using cells with non-targeting sgRNAs. Scatter plots showing probability distributions for gene linkage in cells with control vs targeting sgRNA in sgRNA level analysis. Volcano plot shows discovery pairs that pass significance (p<0.01) as purple dots. d. Single cells clustered at leiden resolution 0.25. Clusters containing similar cell types outlined in specific colors and labeled using respective colors. Individuals and batches were integrated. UMAP visualization of normalized expression of marker genes *HOPX, DCX, DLX5, MKI67, SOX10*, and *NEUROD2*. e. Cell type annotations for each Leiden cluster in the high MOI Perturb-seq experiment, learned through reference mapping to published human developing primary cortex multiomic dataset [61].

**Fig S3:**
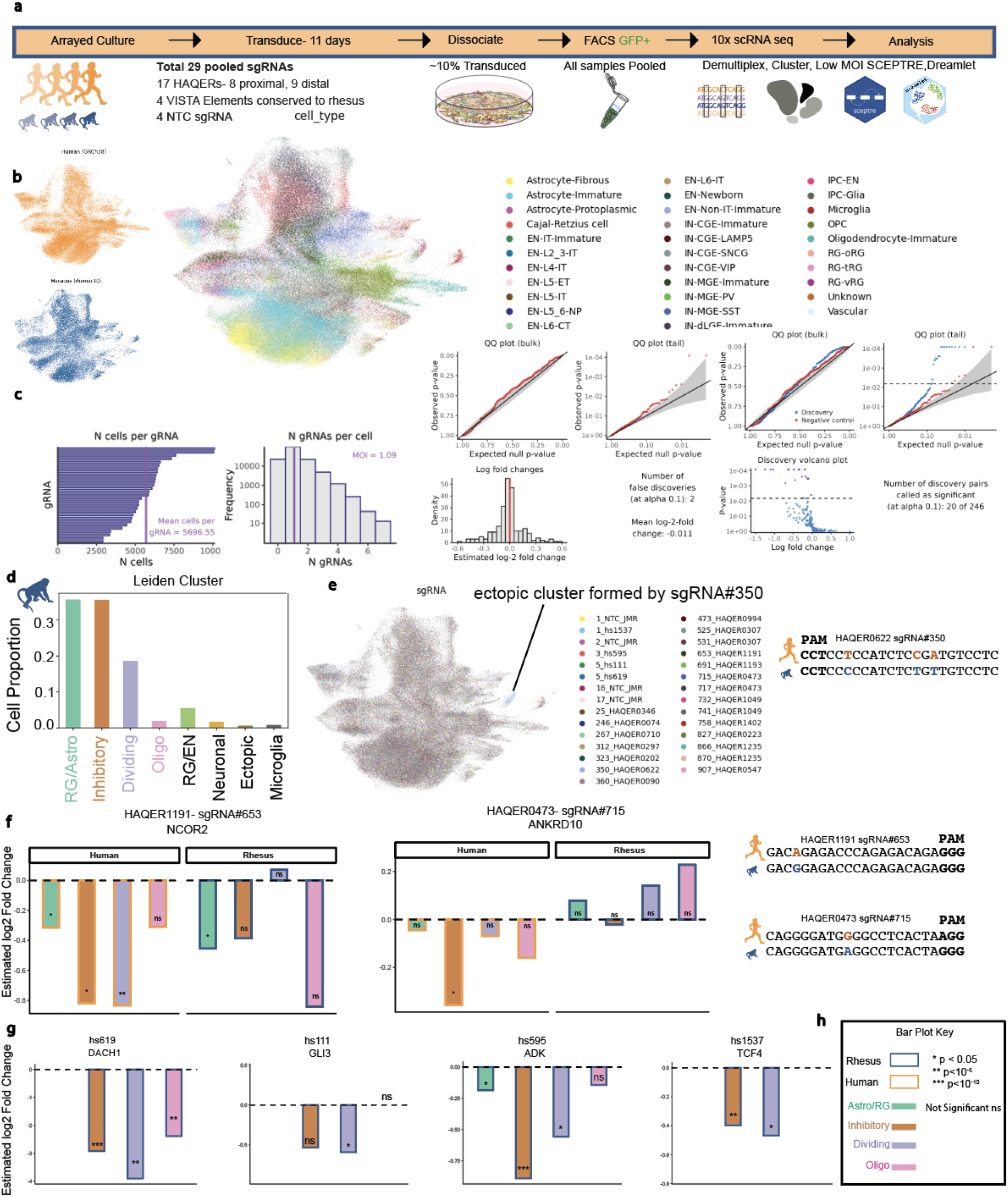
Supplemental analyses for low MOI HAQER Perturb-seq. a. Schematic of the study design including rhesus cells (N=4). Total 29 pooled sgRNAs were used: 17 HAQERs (8 proximal, 9 distal), 4 VISTA elements conserved to rhesus, and 4 non-targeting control (NTC) sgRNAs. Cells were transduced at ∼10% efficiency, pooled, FACS sorted for GFP+, captured by 10x scRNA-seq, and analyzed by demultiplexing, clustering, SCEPTRE, and Dreamlet. b. UMAP of cells from human (orange) and rhesus (blue) colored by cell type annotations assigned after mapping single cells from the low MOI experiment to a published reference dataset [61]. Both species show enrichment for radial glia/astroglia (RG/Astro) as the predominant cell type, consistent with the progenitor-enriching culture conditions. c. Left to right: Histograms of cells per sgRNA and sgRNAs per cell showing MOI ∼1. QQ plots (bulk and tail) during calibration step showing 2 false discoveries at 10% BH FDR at sgRNA level analysis. Histogram shows estimated log2FC for paired genes during calibration step using cells with non-targeting sgRNAs. Volcano plot shows 20 significant regulatory element-gene linkages calculated by sgRNA level analysis. d. Bar plot showing the proportion of cells assigned to each annotated cell type cluster (Leiden clustering, resolution 0.25) in rhesus macaque primary cortical cultures. e. Leiden clustering at resolution 0.25 for species-integrated cells. Ectopic cluster formed by a subset of cells containing sgRNA ID#350 targeting HAQER0622 is labeled on the UMAP (left). sgRNA ID#350 sequence targeting HAQER0622 aligned between human and rhesus (right). PAM site shown in bold. SNPs between human and rhesus are colored in orange or blue respectively. f. Left-Bar plots showing cluster-specific downregulation of genes paired with HAQER-targeting sgRNAs in human vs rhesus cells. HAQER1191-*NCOR2* and HAQER0473-*ANKRD10* linkages are shown across RG/Astro, Inhibitory, Dividing, and Oligo cell clusters for each species. Silencing HAQER1191 paired with significant downregulation of NCOR2 gene in RG/astro, inhibitory, and diving cell types in human samples while in rhesus the significant downregulation was only in the RG/astro cluster. Silencing HAQER0473 showed human-specific ANKRD10 downregulation compared to rhesus (Fig. S3f,h). Conversely, targeting VISTA elements in rhesus led to downregulation of the same paired genes we observed in human samples (Fig. S3g,h). Right-sgRNA sequences targeting HAQERs1191 and 0473 showing 1bp divergence between human and rhesus enabling species comparative differential gene expression. g. Bar plots showing cluster-specific downregulation of genes paired with positive control VISTA elements hs619-*DACH1*, hs111-*GLI3*, hs595-*ADK*, hs1537-*TCF4* in human vs rhesus cells. X axis shows different cell-type clusters and Y axis shows estimated log2 fold change. h. Bar plot color key is shown in the bottom right panel. Significance is indicated as * p<0.05, ** p<10^-5^, *** p<10^-10^; ns = not significant.

**Fig S4:**
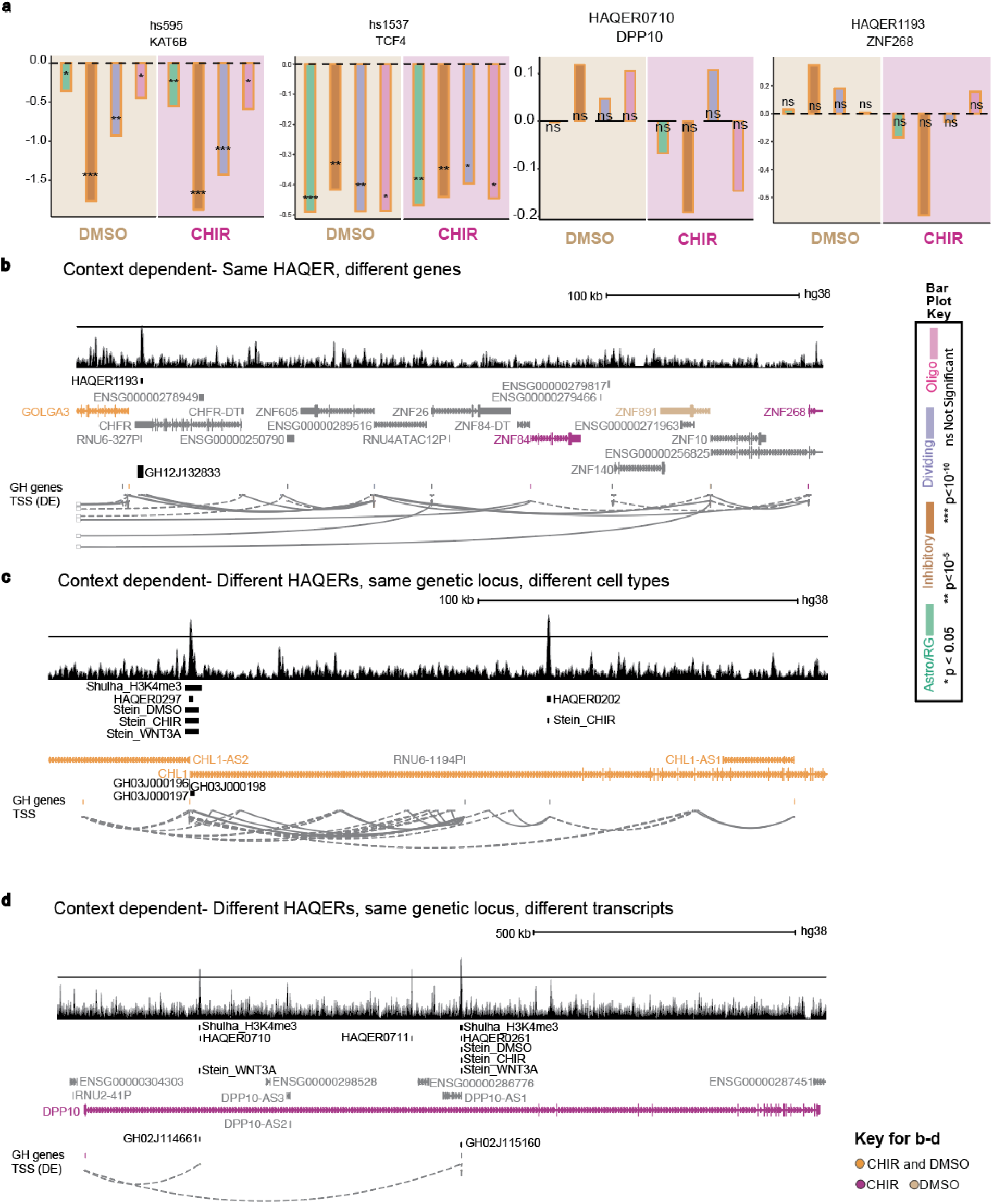
CHIR context dependent regulatory function of HAQERs. a. CHIR context dependent and cell type specific downregulation of genes linked to VISTA elements hs595, hs1537, and non-significant directional trends for HAQER0710 linked to DPP10, and HAQER1193 linked to ZNF268. b. -d. UCSC GeneHancer[26] tracks showing direct and/or indirect looping between HAQERs 1193, 0297, 0202, 0710, 0261 and their nearby genes. Gene bodies are color coded to represent the context in which HAQERs 1193, 0297, and 0710 were significantly paired with the HAQER in the Perturb-seq study (Orange-in both CHIR and DMSO conditions, Magenta-CHIR only, Beige-DMSO baseline only). Tracks beginning with ‘GH’ are GeneHancer tracks, DE notes double elite[26]. Tracks labeled with Shulha and Stein refer to data from the previous studies cited [18,38].

**Fig S5:**
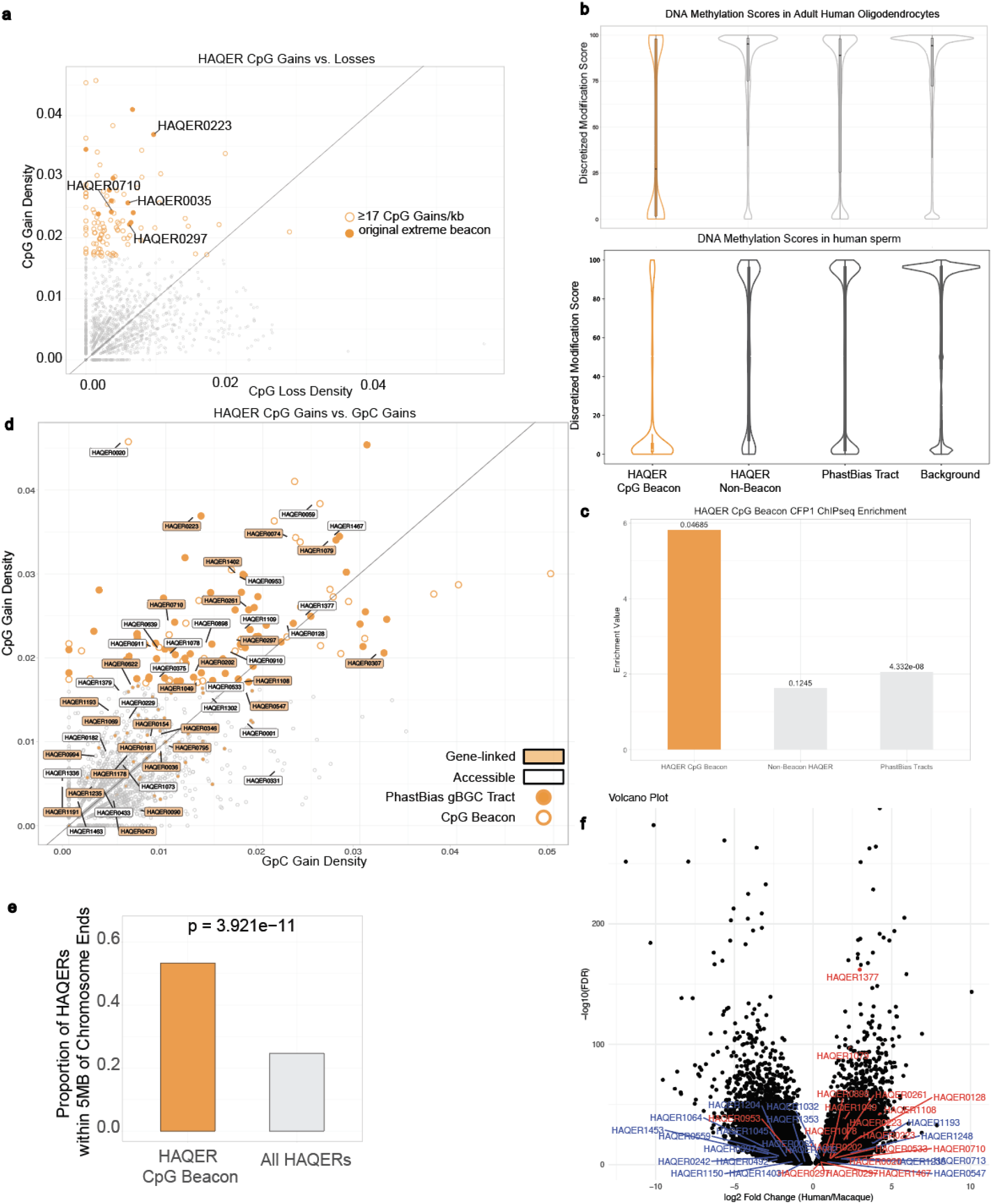
HAQER CpG Beacon Characteristics. a. Scatter plot of CpG gain (Y axis) vs CpG loss (X axis) for all HAQERs. Each dot represents a HAQER. Orange outlined larger circles show HAQER CpG Beacons thresholded by ≥17/kb CpG gain. Note that the 13 previously defined extreme beacon clusters (>20/kb) represent some of the highest densities of CpG gain, labeled in orange filled circles. b. Violin plots of DNA methylation scores of HAQER CpG Beacons (orange), HAQER non-Beacons, phastBias Tracts, and Background genomic regions (all in grey) in adult human oligodendrocytes (top) and human sperm (bottom). HAQER CpG Beacons show significantly reduced DNA methylation compared to all other genomic regions in both cell types (all comparisons significant; exact test statistics in Table S15). c. HAQER CpG Beacons (orange bar), Non-Beacon HAQERs (grey), and phastBias tracts (grey) as groups in the x-axis. Y axis is the enrichment value for CFP1 antibody binding for each of these groups. The numerical value above each bar notes significance, showing HAQER CpG Beacons significantly enriched for CFP1 bound ChIP-seq peaks [83]. d. Scatter plot of HAQER CpG (Y axis) vs GpC gains (X axis). Each dot is a HAQER. HAQER CpG Beacons with >= 17/kb CpG gain have an orange outlined unfilled circle. HAQERs overlapping phastBias regions labeled with orange fill inside the circle. Accessible HAQERs outlined in a black rectangle box. Gene-linked HAQERs, which are a subset of accessible HAQERs, have orange fill inside the rectangle box outlined with black. e. Bar plot showing the proportion of HAQERs within telomeric and subtelomeric regions defined as within 5Mb of chromosome ends (Y axis). X axis shows the 2 groups in comparison, HAQER CpG Beacons vs all HAQERs. f. Volcano plot of orthologous accessible peaks in rhesus and human from bulk ATAC-seq assay under CrossPeak analysis. Each dot is an orthologous ATAC peak. X axis shows the Log2 fold change of accessibility in human/ rhesus. Example peaks overlapping HAQER CpG Beacons labeled in red, non-Beacons in blue. Note that HAQER CpG Beacons show increased accessibility in human samples compared to non-Beacon HAQERs.

## Supplementary Tables

1. Primary human and rhesus macaque sample info-Age and ID, 10x scRNA-seq samples, bulk ATAC-seq samples
2. Human (hg38 coordinates) and rhesus accessible HAQERs
3. Twist sgRNA oligo order with overhangs for cloning
4. High MOI gRNA-target input table for SCEPTRE
5. High MOI SCEPTRE element level analysis output table
6. High MOI SCEPTRE sgRNA level analysis output table
7. Low MOI gRNA-target input table for SCEPTRE
8. Low MOI Validation SCEPTRE output sgRNA level output table
9. Low MOI DMSO baseline SCEPTRE output sgRNA level output table
10. Low MOI CHIR stimulation SCEPTRE output sgRNA level output table
11. HAQER regioneR[72] overlaps with H3K4me3[38] and CpG[39,42] -hg19
12. HAQERs overlapping with Extreme CpG Beacon Clusters from Bell et al 2014 -hg19
13. HAQER CpG Beacons defined from our study (>= 17/kb CpG)
14. HAQERs overlapping the top 10 regions of gBGC from Dreszer et al 2007 -hg18
15. DNA Methylation test statistics in adult human neuron, oligodendrocyte, and sperm

